# Serotonin sensing by microglia conditions the proper development of neuronal circuits and of social and adaptive skills

**DOI:** 10.1101/2022.03.09.483609

**Authors:** Giulia Albertini, Ivana D’Andrea, Mélanie Druart, Catherine Béchade, Nayadoleni Nieves-Rivera, Fanny Etienne, Corentin Le Magueresse, Alexandra Rebsam, Nicolas Heck, Luc Maroteaux, Anne Roumier

## Abstract

The proper maturation of emotional and sensory circuits requires a fine tuning of serotonin (5-HT) level during early postnatal development. Consistently, dysfunctions of the serotonergic system have been associated with neurodevelopmental psychiatric diseases, including autism spectrum disorders (ASD). However, the mechanisms underlying the developmental effects of 5-HT remain partially unknown, one obstacle being the action of 5-HT on different cell types.

Here, we focused on microglia, which play a role in brain wiring refinement, and we investigated whether the control of these cells by 5-HT is relevant for neurodevelopment and spontaneous behaviors. Since the main 5-HT sensor in microglia is the 5-HT_2B_ receptor subtype, we prevented 5-HT signaling specifically in microglia by conditionally invalidating *Htr2b* gene in these cells. We observed that abrogating the serotonergic control of microglia neonatally impacts the phagolysosomal compartment of these cells and their proximity to synapses, and perturbs neuronal circuits maturation. Furthermore, this early ablation of microglial 5-HT_2B_ receptors leads to adult hyperactivity in a novel environment and behavioral defects in sociability and flexibility. Importantly, we show that these behavioral alterations result from a developmental effect, since they are not observed when microglial *Htr2b* invalidation is induced later, at P30 onward.

Thus, a primary alteration of 5-HT sensing in microglia, during a critical time window between birth and P30, is sufficient to impair social and flexibility skills. This link between 5-HT and microglia may explain the association between serotonergic dysfunctions and behavioral traits, like impaired sociability and inadaptability to novelty, which are prominent in several psychiatric disorders such as ASD.

## Introduction

Neurodevelopmental psychiatric disorders such as autism spectrum disorders (ASD) are characterized by alterations of spontaneous behaviors and personality traits affecting notably social interactions and adaptability to novelty. Despite their high prevalence - roughly one percent of the total population for ASD -, preventive, diagnosis and therapeutic actions for these disorders are still limited by lack of understanding of their underlying pathophysiology. Among potential risk factors for neurodevelopmental psychiatric disorders are events or mutations that affect the immune system and notably microglia, the brain resident macrophages, which participate to neuronal migration, synapse formation and elimination^1^. An imbalance of the neuromodulator serotonin (5-HT)^2–4^ has also been incriminated. Indeed, hyperserotonemia was one of the first biomarkers identified for autism^5^, and is observed in 25% of ASD patients with current criteria. Furthermore, 5-HT has developmental roles^6–8^, and in mice, increasing or decreasing 5-HT availability during development, through genetic or pharmacological means, alters emotional and sensory neuronal circuits maturation, and has long-term effects on anxiety and sociability in adulthood^6–8^

The mechanisms underlying the implication of 5-HT in the etiology of neurodevelopmental disorders remain partly elusive due to the potential action of 5-HT on different cell types. Notably, besides neurons, which express a large array of serotonergic receptors, microglia express the 5-HT receptor subtype 2B (5-HT_2B_) throughout postnatal life^9–11^ and we have shown that 5-HT triggers a directional motility, 5-HT_2B_-dependent, of microglial processes^9,12^. In addition, *Htr2b^-/-^* mice, fully invalidated for the gene encoding 5-HT_2B_ receptors, show defects in the maturation of the retinal circuit in the thalamus^9^, a developmental process of synapse elimination requiring both microglia and 5-HT^8,13–15^, and a wide spectrum of behavioral alterations^16^. However, the interpretation of these *Htr2b^-/-^* mice phenotypes is limited by the fact that *Htr2b* gene is also expressed by subsets of serotonergic and dopaminergic neurons^17,18^. Considering the prominent role of microglia in sculpting brain circuits, and their ability to respond to 5-HT, we tested here the hypothesis that impaired 5-HT sensing specifically in microglia is sufficient to alter the refinement of postnatal circuits, and thereby to induce alterations in spontaneous behaviors relevant to neurodevelopmental psychiatric disorders.

To this aim, we invalidated *Htr2b* gene specifically in the microglia/macrophage lineage early in postnatal development, by using *Cx3cr1^creERT2/+^;Htr2b^fl/fl^* mice treated with tamoxifen just after their birth, named hereafter cKO^Htr2b-μglia^ _TXFbirth_. The controls were *Cx3cr1^creERT2/+^; Htr2b^+/+^* mice similarly treated with tamoxifen, and named cWT_TXFbirth_ hereafter. Importantly, to disentangle possible developmental vs. adult effects of 5-HT signaling in microglia, we also performed experiments with mice of the same genotypes treated with tamoxifen after four weeks of postnatal development, a time point hereafter referred to as “P30”. These mice are named cKO^Htr2b-μglia^ _TXF-P30_ and cWT_TXF-P30_. Our results show that impairing 5-HT signaling in microglia since birth perturbs microglial development and the refinement of neuronal circuits, and leads to hyperactivity in a novel environment, poor sociability and limited behavioral flexibility. Importantly, such behavioral alterations are not induced by ablation of the microglial 5-HT_2B_ receptors after P30, demonstrating a developmental role of the serotonergic regulation of microglia for these spontaneous behaviors. Altogether, our results support the idea that 5-HT, via the 5-HT_2B_ receptor, controls microglia and some of their well-established effects on brain development, and that an early alteration of this serotonergic control can contribute to the etiology of neurodevelopmental disorders characterized by sociability and flexibility defects, such as ASD.

## Results

### 1. Absence of 5-HT_2B_ receptors in neonatal microglia decreases their lysosome content, alters their morphology, and impacts the synapse-microglia proximity in the developing brain

During the first weeks of postnatal development, microglia proliferate and undergo morphological and functional maturation, with changes in their phagocytic activity^19,20^. A precise timing of these changes is necessary for microglia to properly regulate brain circuits formation, through the induction of dendritic filopodia formation^21^, engulfment of presynaptic elements^13,22,23^, or promotion of spine maturation^24^, but the determinants of this maturation are unknown. To investigate whether microglia development was impacted by lack of serotonergic sensing we triggered recombination of *Htr2b* in microglia with tamoxifen administration between postnatal day (P)1 and P5 (“TXF birth”)^11^ and characterized microglia at P15 (timeline in **Fig. 1a**). Data from males and females were pooled except for parameters when a sex-effect was observed (e.g. for morphology, see hereafter). First, we checked the microglial (Iba1^+^ cells) density and did not detect effect of the genotype in any of the brain regions we analyzed (hippocampus, cortex, dorsal lateral geniculate nucleus (dLGN) of the thalamus) (**Fig. S1a-b**). Then, as developing microglia normally exhibit high phagocytic activity, which has been linked to synaptic pruning and refinement of brain connectivity^22,25,26^, we measured the percentage of microglial (Iba1^+^) volume occupied by CD68, a phagolysosomal marker upregulated in actively phagocytic cells^27^. This revealed that the lysosomal content was strongly reduced (−40%) in hippocampal microglia of P15 cKO^Htr2b-μglia^_TXFbirth_ compared to cWT_TXFbirth_ mice (**Fig. 1b-c**). During postnatal development, microglia also undergo a transition from amoeboid to ramified ^19,20^, in a sex- and region-dependent manner^28,29^. We thus performed three-dimensional (3D) reconstructions of individual CA1 microglia at P15 (examples in **Fig. 1d**) to assess their morphology. In male mice, 3D Sholl analysis showed a reduced complexity of microglial processes in cKO^Htr2b-μglia^_TXFbirth_ compared to cWT_TXFbirth_ (**Fig. 1e**), as well as a significant decrease in total length of processes and the number of processes terminal points (**Fig. 1f-g**). Noteworthy, these differences were not detected in females (**Fig. 1h-j**), indicating a sex-dependent effect of 5-HT_2B_-mediated signaling on microglia morphology.

**Fig. 1.**
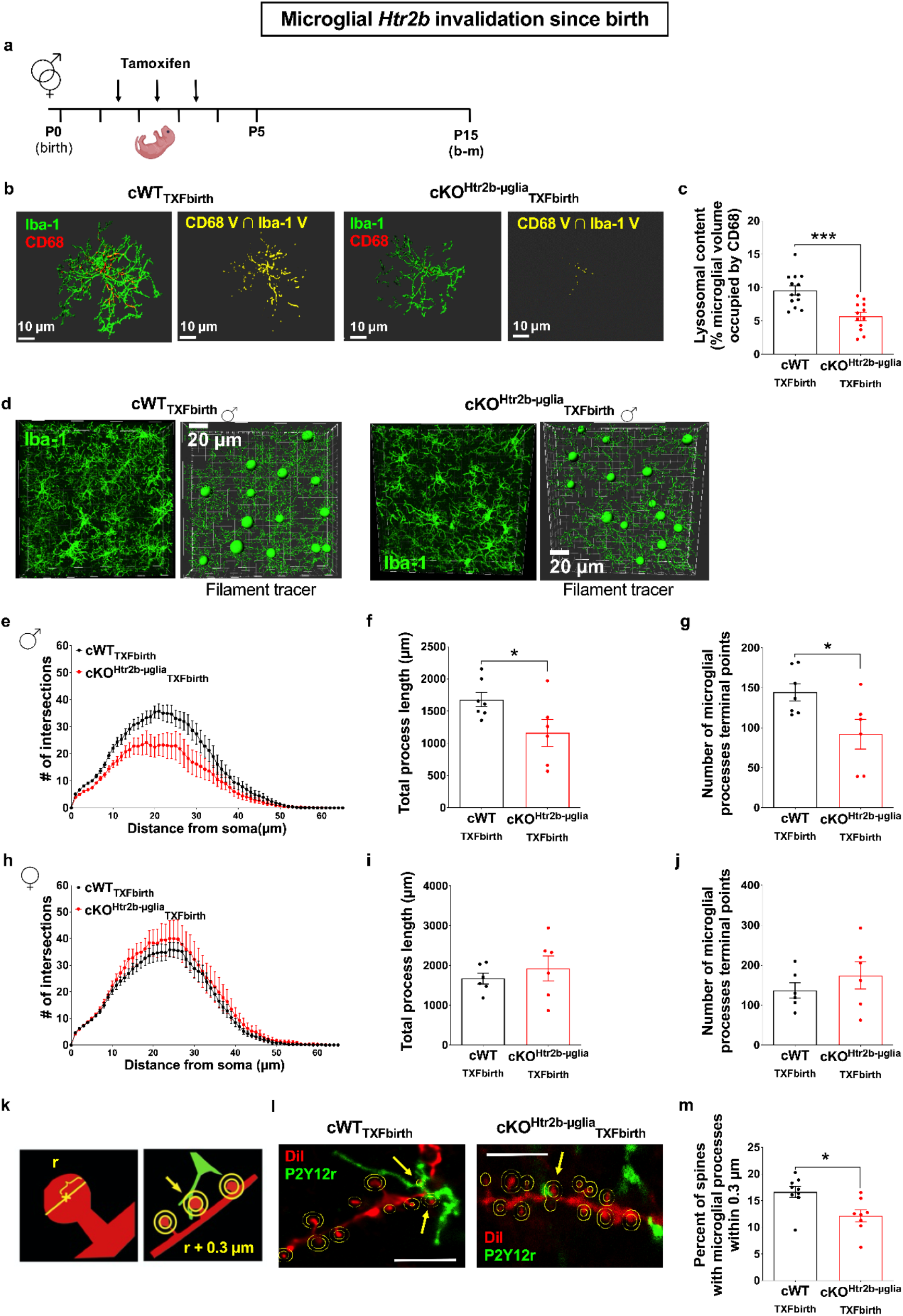
Absence of 5-HT_2B_ receptors in neonatal microglia decreases their lysosome content, alters their morphology, and impacts the synapse-microglia proximity in the developing brain. **a**, Timeline of tamoxifen treatment for early *Htr2b* invalidation in microglia, and the analyses performed at P15. **b**, Representative images of CA1 microglia and lysosomal content in cWT_TXFbirth_ and cKO^Htr2b-μglia^_TXFbirth_ mice at P15. Left panels: immunostaining for Iba1 and microglial lysosome membrane protein CD68; right panels: 3D reconstruction of CD68^+^ lysosomes with Iba1^+^ cells overlay (in yellow). **c,** The relative microglial cell volume occupied by CD68^+^ lysosomes is strongly reduced in cKO^Htr2b-μglia^_TXFbirth_ mice at P15, as compared to cWT_TXFbirth_ mice (cWT_TXFbirth_ mice, *n* = 13 mice; cKO^Htr2b-μglia^_TXFbirth_ mice, *n* = 12 mice; 8 microglia were analyzed and averaged per animal; unpaired t-test with Welsh’s correction, p = 0.0003). **d**, Representative images of CA1 Iba1^+^ cells (left panels) and 3D reconstructions carried out using the filament tracer module by Imaris (right panels), here in male mice at P15. **e**, Sholl profiles of microglia in male mice at P15; note the reduced arborization in cKO^Htr2b-μglia^_TXFbirth_ (two-way RM-ANOVA: significant main effects of interaction P < 0.0001, F_65,715_ = 2.571, radius P < 0.0001, F_65,715_ = 65.13 and genotype P = 0.0295, F_1,11_ = 6.254). **f**, Total process length of Iba1^+^ cells ramifications in male mice at P15; note the reduced total length of microglial processes in cKO^Htr2b-μglia^_TXFbirth_ (Mann-Whitney test, p = 0.0350). **g**, Number of microglial processes terminal points in male mice at P15; note the reduced number of terminal points in cKO^Htr2b-μglia^_TXFbirth_ (Mann-Whitney test, p = 0.0408). **h**, Similar Sholl profiles of microglia in cWT_TXFbirth_ and cKO^Htr2b-μglia^_TXFbirth_ female mice at P15. **i**, Comparable total process length of Iba1^+^ cells ramifications in cWT_TXFbirth_ and cKO^Htr2b-μglia^_TXFbirth_ female mice at P15. **j**, Similar number of microglial processes terminal points in cWT_TXFbirth_ and cKO^Htr2b-μglia^_TXFbirth_ female mice at P15. **k,** Drawings of the method used for quantifying potential association of microglial processes with dendritic spines. **l,** Representative images of hippocampal dendrites of cWT_TXFbirth_ and cKO^Htr2b-μglia^_TXFbirth_ mice surrounded by P2Y12 receptor (P2Y12r)^+^ microglial processes at P15. The arrows indicate the spines with a microglial process in close proximity (< 0.3 μm). Analysis was performed in 3D in image stacks, the figure is represented in 2D for easier visualization. Scale bar: 5 μm. **m**, The percent of spines located within 0.3 μm to microglial processes was significantly reduced in cKO^Htr2b-μglia^_TXFbirth_ mice (cWT_TXFbirth_ mice, *n* = 9 mice; ecKO^Htr2b-μglia^_TXFbirth_ mice, *n* = 8 mice; Mann-Whitney test, P = 0.0111). **e-j**: cWT_TXFbirth_ males, *n* = 7 mice; cKO^Htr2b-μglia^_TXFbirth_ males, *n* = 6 mice; cWT_TXFbirth_ females, *n* = 6 mice; cKO^Htr2b-μglia^_TXFbirth_ females, *n* = 6 mice. For each mouse, the value represents the average value of 4-8 microglia. Graphs show mean±s.e.m. and points represent individual animals. *p< 0.05, ***p < 0.005.

Physical contacts between microglia and dendritic spines participate to the sculpting of postnatal neuronal networks^21^. We thus analyzed the structural association of microglial processes with hippocampal dendritic spines. To do so, we performed DiI and P2Y12 receptor labeling to simultaneously visualize CA1 apical dendrites and microglia, and we measured the percentage of dendritic spines with microglial processes located within 0.3 μm of the spine head (scheme in **Fig. 1k** and representative images in **Fig. 1l**). By this approach, we determined that the proportion of spines in close proximity to microglia was reduced (by 30%) in cKO^Htr2b-μglia^_TXFbirth_ mice (**Fig. 1m**). Of note, the probability for a spine to have a microglial process in its surroundings increases with its head diameter, and thus spine head size can be a confounding factor, but we verified that the proportion of spines apposed to a microglial process was reduced in cKO^Htr2b-μglia^_TXFbirth_ mice whatever the spine head size (**Fig. S1c**). As this effect was observed despite unchanged microglia density, and both in males and females whereas the latter have a normal microglia morphology, it may indicate an intrinsic reduced ability of microglia to structurally interact with spines in the absence of 5-HT_2B_ receptor signaling.

In summary, ablation of 5-HT_2B_ receptor in microglia since birth perturbs microglial maturation and the contacts of dendritic spines during early postnatal development.

### 2. Absence of 5-HT_2B_ receptors in microglia since birth impairs synaptic and axonal refinement in the developing hippocampus, cortex and thalamus, respectively

The first weeks of postnatal development are marked by intense synapse formation and pruning of connectivity, which preclude the establishment of mature neuronal circuits. Having identified functional and structural defects in P15 microglia when their 5-HT sensing was disturbed, we investigated whether this may have impacted connectivity in cKO^Htr2b-μglia^_TXFbirth_ mice at the same age. We focused on hippocampus, cortex and thalamus, i.e. regions where microglia have been involved in developmental axonal or synaptic refinement^13,23–26,30^. Data from males and females were pooled after checking for the absence of sex-dependent effect. In the hippocampus, we observed with Golgi staining (**Fig. 2a**) that both the density and the length of dendritic protrusions (spines and filopodia-like spines) were increased in CA1 pyramidal neurons of cKO^Htr2b-μglia^_TXFbirth_ compared to cWT_TXFbirth_ mice (**Fig. 2b, d**). Accordingly, the distribution of protrusion types, as defined in^31^, was significantly altered in cKO^Htr2b-μglia^_TXFbirth_ mice (**Fig. 2e**). Of note, no differences in the spine head diameter was observed (**Fig. 2c**), further confirming no spine head-related bias in the previous spine-microglia proximity phenotype (**Fig. 1f**). Interestingly, similar increases in protrusions density and length, and alteration of protrusion types distribution, were observed in the prefrontal cortex (L2/L3 principal neurons) (**Fig. S2a-e**). A functional analysis, performed by whole-cell recording of CA1 pyramidal neurons on acute brain slices (**Fig. 2f**), confirmed the existence of synaptic alterations. Indeed, hippocampal neurons from cKO^Htr2b-μglia^_TXFbirth_ P15 mice displayed higher mEPSC frequency compared to controls (**Fig. 2g**), despite no changes in amplitude (**Fig. 2h**). Thus, the early impairment of 5-HT sensing in microglia induces synaptic alterations in the hippocampus and cortex, which are consistent with impaired synaptic pruning and/or maturation.

**Fig. 2.**
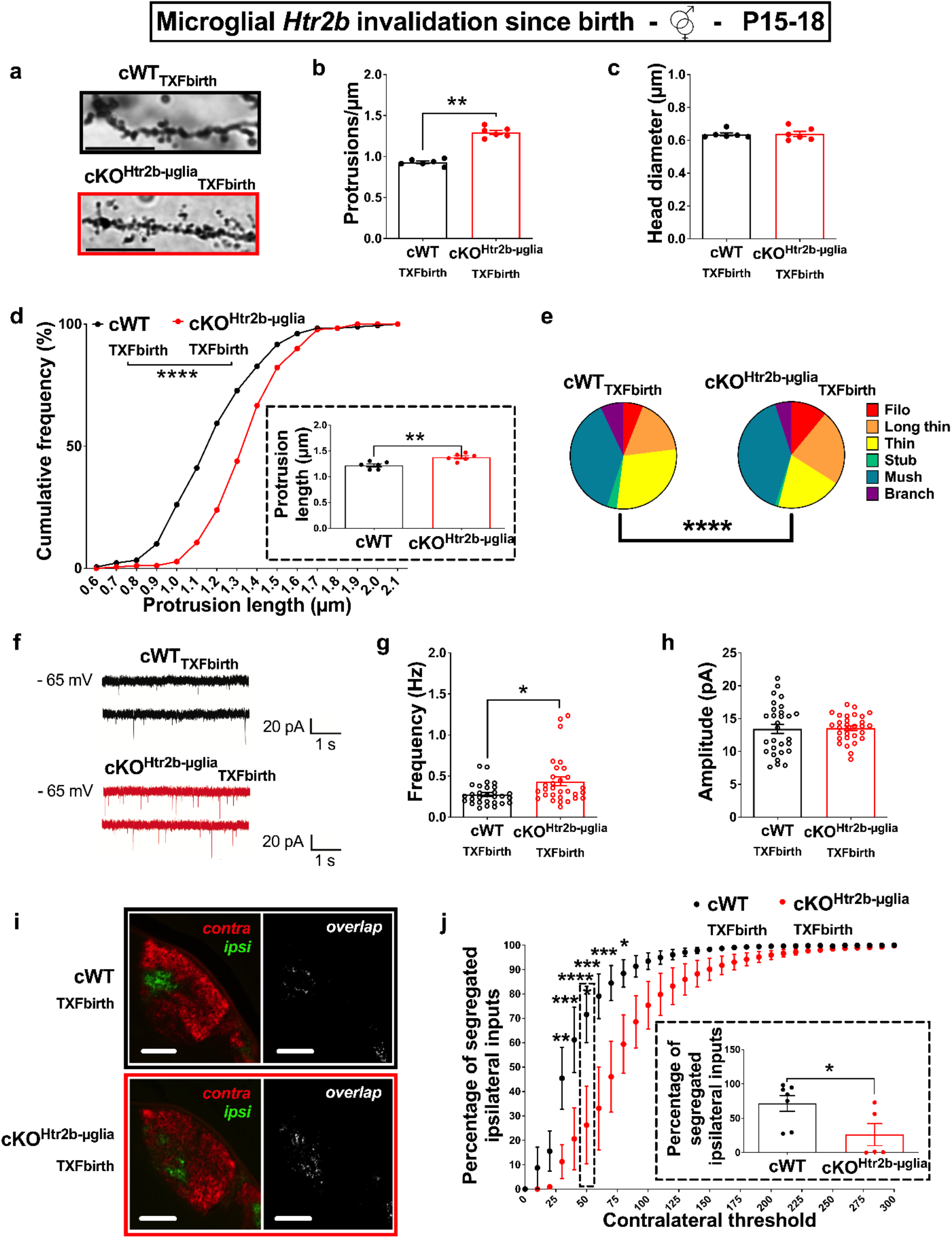
Absence of 5-HT_2B_ receptors in microglia since birth impairs synaptic and axonal refinement in the developing hippocampus and thalamus, respectively. **a,** Representative images of secondary apical dendrites in hippocampal principal neurons labeled with Golgi staining, scale bar: 10 μm. **b**, Quantification of protrusion density on secondary apical dendrites of CA1 principal neurons in cWT_TXFbirth_ and cKO^Htr2b-μglia^_TXFbirth_ mice at P15 obtained averaging 30 dendrites/mouse, that shows a significant increase in cKO^Htr2b-μglia^_TXFbirth_ mice (Mann-Whitney test, p = 0.0022). **c**, Similar average head diameter of protrusions on secondary apical dendrites of CA1 principal neurons in cWT_TXFbirth_ and cKO^Htr2b-μglia^_TXFbirth_ mice at P15. **d**, Cumulative distribution plot for the average length of protrusions on secondary apical dendrites of CA1 principal neurons in cWT_TXFbirth_ and cKO^Htr2b-μglia^_TXFbirth_ mice at P15; note the significant rightward shift in cKO^Htr2b-μglia^_TXFbirth_ mice (Kolmogorov-Smirnov test, p < 0.0001). In the inset, quantification of protrusion length obtained averaging 30 dendrites/mouse, that confirms a significant increase in cKO^Htr2b-μglia^_TXFbirth_ mice (Mann-Whitney test, p = 0.0087). **e**, Pie charts showing the altered distribution of protrusion types in cKO^Htr2b-μglia^_TXFbirth_ mice, as compared to cWT_TXFbirth_ mice at P15 (Chi-square test, p < 0.0001). Abbreviations: Filo: filopodia; Stub: stubby; Mush: mushroom. **f**, Representative traces of mEPSC recordings in CA1 hippocampal acute slices from cWT_TXFbirth_ and cKO^Htr2b-μglia^_TXFbirth_ mice at P15. **g**, Increased mESPC frequency in cKO^Htr2b-μglia^_TXFbirth_ mice at P15, as compared to cWT_TXFbirth_ mice (Mann-Whitney test, p = 0.0120). **h**, Similar mESPC amplitude in cWT_TXFbirth_ and cKO^Htr2b-μglia^_TXFbirth_ mice at P15. **i**, Dye-labeled retinogeniculate projections in dLGN of cWT_TXFbirth_ and cKO^Htr2b-μglia^_TXFbirth_ mice at P17-P18, 48h after intravitreal injection of AlexaFluor 488 or 555 in either eye. Left panels: projections from retinal ganglion cells from the ipsilateral retina are in green and those from the contralateral retina are in red; right panels: overlap between ipsi- and contralateral projections. Scale bars 100 μm. **j**, Percentage of pixels containing only ipsilateral signal (no contralateral signal) as a function of contralateral threshold (ipsilateral threshold is fixed) in dLGN of P17-18 cWT_TXFbirth_ and cKO^Htr2b-μglia^_TXFbirth_ mice (*n* = 7 and 5 mice, respectively); note the significant rightward shift in cKO^Htr2b-μglia^_TXFbirth_ mice, that demonstrate that ipsilateral fibers are less segregated from contralateral fibers (thus more overlapped) than in cWT_TXFbirth_ mice (two-way RM-ANOVA: significant main effects of interaction P < 0.0001, F_30,300_ = 4.118, contralateral threshold P = 0.0288, F_30,300_ = 76.03 and genotype P = 0.0295, F_1,10_ = 6.507); Sidak multiple comparisons: 30, p = 0.0027; 40, p = 0.0001; 50, p < 0.0001; 60, p < 0.0001; 70, p = 0.0003; 80, p = 0.0256. The inset shows a columnar representation of the values for the contralateral threshold set at 50 (Mann-Whitney test, p = 0.0303). **Fig. b-c, d (inset), e**, *n* = 6 mice/genotype, average of 30 dendrites per mouse; **Fig. d**, *n* = 180 dendrites from 6 mice/genotype; **Fig. g-h**, cWT_TXFbirth_ mice, *n* = 29 cells from 5 mice; cKO^Htr2b-μglia^_TXFbirth_ mice, *n* = 30 cells from 5 mice. Graphs show mean±s.e.m. and points represent individual animals. *p < 0.05, **p < 0.01, ***p < 0.005, ****p < 0.0001.

Another region undergoing prominent refinement soon after birth is the dLGN of the thalamus^13,22,23^, where eye-specific segregation of retinal projections requires both appropriate 5-HT levels^32^ and functional microglia^13^. We had previously demonstrated that the maturation of this visual circuit was impaired in *Htr2b* full knock-out, mice^9^. Here, the conditional knock-out model allowed us to test if the axonal refinement in the dLGN was dependent on the microglial 5-HT_2B_ receptor. We observed that ipsilateral and contralateral retinal inputs were actually significantly less segregated, i.e. more overlapped, in cKO^Htr2b-μglia^_TXFbirth_ than in cWT_TXFbirth_ mice at P17-18, a time point when segregation is normally achieved (**Fig. 2i-j**). Such defective axonal refinement was confirmed by the fact that ipsilateral projections in cKO^Htr2b-μglia^_TXFbirth_ mice were scattered over a greater dLGN area than in cWT_TXFbirth_ mice (**Fig. S2f-h**).

Overall, our results in different brain areas demonstrate that impairment of 5-HT sensing in microglia since birth affects synaptic pruning, axonal refinement and thus brain wiring in the first weeks of postnatal development.

### 3. Absence of 5-HT_2B_ receptors in microglia since birth impacts activity of mice in a novel environment, sociability and flexibility

Alterations of brain development can be either compensated with age or on the contrary lead to persistent behavioral deficits. To assess long-lasting consequences of interrupting 5-HT_2B_ receptor-mediated signaling in microglia since birth, we performed a battery of tests to profile the behavior of adult (or, for one test, juvenile) cKO^Htr2b-μglia^_TXFbirth_ mice in three domains commonly altered in neurodevelopmental psychiatric disorders: (i) behavior in a novel environment; (ii) social skills; (iii) flexibility (time line in **Fig. 3a**). All behavioral experiments were performed and analyzed separately on males and females, with the most significant alterations being observed in males. Indeed, when adult cKO^Htr2b-μglia^_TXFbirth_ male mice were exposed to a novel environment, they responded with significantly increased locomotion (**Fig. 3b-c**) compared to cWT_TXFbirth_ mice. This effect seemed to be due to the novelty of the environment rather than hyperactivity per se, being more prominent at the beginning of the test. Moreover, when placed in an unfamiliar cylinder, male cKO^Htr2b-μglia^_TXFbirth_ mice spent significantly more time self-grooming than cWT_TXFbirth_ mice, suggesting an increased propensity to repetitive stereotypic behavior under unknown and stressful conditions (**Fig. 3d-e**).

**Fig. 3.**
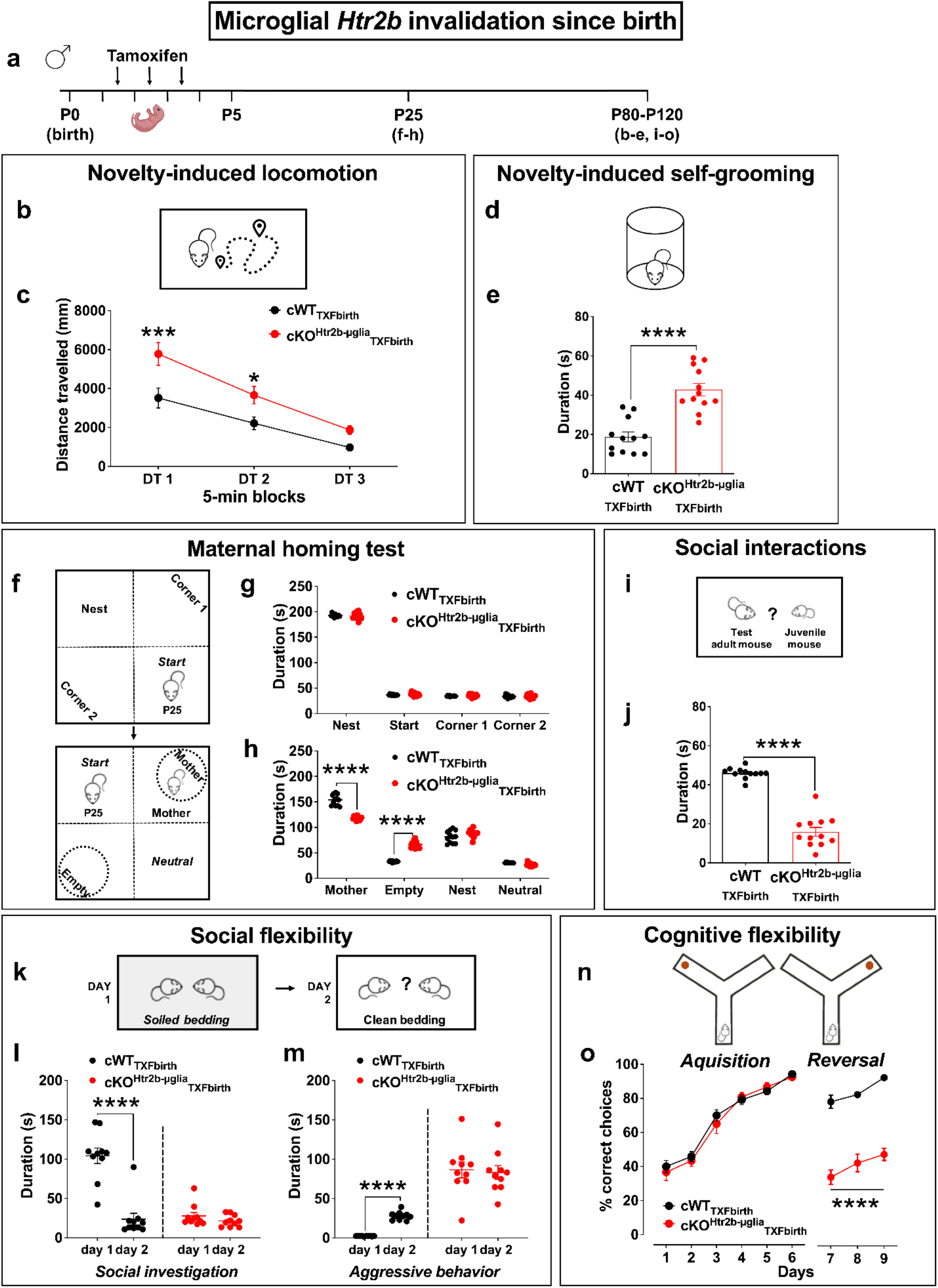
Absence of 5-HT_2B_ receptors in microglia since birth impacts activity in a novel environment, sociability and flexibility in male mice. **a,** Timeline of tamoxifen treatment for early microglial *Htr2b* invalidation, and the behavioral tests performed in juvenile (maternal homing test) or adult male mice. **b**, Scheme of the approach used to assess locomotion, in a novel arena. **c**, Distance travelled in response to a novel environment in 5-min blocks (DT) (two-way RM-ANOVA: significant main effect of DT P < 0.0001, F2,44 = 69.85 and genotype P = 0.0040, F1,22 = 10.36); note that the increased locomotion is significant only in the first 10 min of test (Sidak multiple comparisons: cWT_TXFbirth_ vs. cKO^Htr2b-μglia^_TXFbirth_, DT1, p = 0.0006; DT2, p = 0.0416), suggesting that this effect is due to the novelty of the environment rather than to hyperactivity per se. **d**, Scheme of the approach used to assess self-grooming, in an unknown cylindrical arena. **e**, The time spent grooming in response to a novel environment is strongly enhanced in cKO^Htr2b-μglia^_TXFbirth_ compared to cWT_TXFbirth_ adult male mice (unpaired *t*-test, p < 0.0001). **f**, Scheme of maternal homing test: habituation (upper panel) and test (lower panel). **g,** Similar behavior during the habituation period by cWT_TXFbirth_ and cKO^Htr2b-μglia^_TXFbirth_ P25 male mice. **h**, Time spent in each corner during the homing test (two-way RM-ANOVA: significant main effects of interaction *P* < 0.0001, *F*_3,66_ = 77.11, zone *P* < 0.0001, *F*_3,66_ = 847.0 and genotype *P* = 0.0004, *F*_1,22_ = 17.74); note that the preference for the mother corner was strongly reduced in cKO^Htr2b-μglia^_TXFbirth_ P25 male mice (Sidak multiple comparisons: mother corner, cWT_TXFbirth_ vs. cKO^Htr2b-μglia^_TXFbirth_, p < 0.0001) and, conversely, we observed an increased preference for the empty corner in cKO^Htr2b-μglia^_TXFbirth_ mice (Sidak multiple comparisons: empty corner, cWT_TXFbirth_ vs. cKO^Htr2b-μglia^_TXFbirth_, p < 0.0001). **i**, Scheme of the approach used to assess social interactions between test mice and unfamiliar juveniles performed in a neutral territory. **j**, The time spent interacting with a juvenile is strongly reduced in cKO^Htr2b-μglia^_TXFbirth_ adult male mice (unpaired *t*-test, p < 0.0001). **k**, Scheme of the approach used to assess spontaneous social behavior in the home cage and social flexibility of adult male mice, using the change of litter bedding on day 2 as an environmental challenge. **l**, Duration of social investigation, indicative of affiliative behavior (twoway RM-ANOVA: significant main effects of interaction *P* < 0.0001, *F*_1,18_ = 64.13; day *P* < 0.0001, *F*_1,18_ = 87.52 and genotype *P* = 0.0098, *F*_1,18_ = 22.76); note that only cWT_TXFbirth_ male mice modified their behavior in agreement to the change in social hierarchy induced by environmental challenge, showing higher duration of social investigation on day 1 (Sidak multiple comparisons: cWT_TXFbirth_, day 1 vs. day 2, p < 0.0001), when the social hierarchy was defined, compared to day 2, when the social hierarchy needed to be established. **m**, Duration of aggressive behavior (twoway RM-ANOVA: significant main effects of interaction *P* < 0.0001, *F*_1,18_ = 68.52; day *P* < 0.0001, *F*_1,18_ = 40.81 and genotype *P* < 0.0001, *F*_1,18_ = 55.91); note that, on day 2, only cWT_TXFbirth_ male mice showed an increased aggressive behavior compared to day 1 (Sidak multiple comparisons: cWT_TXFbirth_, day 1 vs. day 2, p < 0.0001). **n**, Scheme of the Y-maze reversal learning task. In the acquisition phase, mice were trained to find the food reward in one arm of the Y-maze. In the reversal procedure, the reward location was switched to the arm opposite to its previous location. **o**, During the acquisition phase, all mice reached the criterion to choose the correct arm of the Y-maze; however, cKO^Htr2b-μglia^_TXFbirth_ male mice failed to perform the reversal phase of the test (twoway RM-ANOVA: significant main effects of phase *P* = 0.0001, *F*_2,44_ = 11.28 and genotype *P* < 0.0001, *F*_1,22_ = 138.0); Sidak multiple comparisons: cWT_TXFbirth_ vs. cKO^Htr2b-μglia^_TXFbirth_, day 7, p < 0.0001; day 8, p < 0.0001; day 9, p < 0.0001. **c, e, g-h, j, o**: cWT_TXFbirth_, *n* = 12 male mice; cKO^Htr2b-μglia^_TXFbirth_, *n* = 12 male mice. **l-m**: cWT_TXFbirth_, *n* = 10 male mice; cKO^Htr2b-μglia^_TXFbirth_, *n* = 10 male mice. Graphs show mean±s.e.m. and points represent individual animals. *p < 0.05, ***p < 0.005, ****p < 0.0001.

To identify early deficits in social behavior, we performed the maternal homing test on juvenile mice (P25, **Fig. 3f**). In the first configuration of the test, juvenile cKO^Htr2b-μglia^_TXFbirth_ male mice spent significantly more time in the corner with home cage bedding than with fresh bedding (**Fig. 3g**), demonstrating an intact ability to recognize familiar olfactory cues. However, in the second configuration where we introduced two mesh tubes, one empty and one containing the juvenile’s mother, the time spent in the corner containing the latter tube was significantly reduced, and the time spent in the corner containing the empty tube was increased, in cKO^Htr2b-μglia^_TXFbirth_ mice compared to cWT_TXFbirth_ (**Fig. 3h**). The social deficit observed in cKO^Htr2b-μglia^_TXFbirth_ juvenile mice toward their mother persisted in adulthood toward unfamiliar mice. Indeed, in a neutral territory (**Fig. 3i**), cKO^Htr2b-μglia^_TXFbirth_ adult male mice spent less time interacting with an unfamiliar (wild-type) juvenile compared to cWT_TXFbirth_ (**Fig. 3j**). We further explored the social skills of male mice by investigating their social flexibility. To this aim, we recorded the spontaneous social behavior in home cage of male mice housed with a familiar cage mate, during two consecutive days, using the change of litter bedding after day 1 to challenge the social hierarchy (**Fig. 3k**)^33^. As expected, cWT_TXFbirth_ mice adapted their behavior: on day 2, they showed less social investigation (**Fig. 3l**) and more aggressive behavior (**Fig. 3m**) than on day 1 when social hierarchies were defined. By contrast, cKO^Htr2b-μglia^_TXFbirth_ mice spent less time interacting with the conspecific (**Fig. 3l**) and showed an abnormal aggressive behavior overall (**Fig. 3m**), independently of the environmental challenge. These results confirm a strong social impairment and reveal traits of social inflexibility in cKO^Htr2b-μglia^_TXFbirth_ mice. To understand if the impairment in flexibility could be generalized beyond the social domain, we assessed cognitive flexibility with a reversal learning test (**Fig. 3n**). During the acquisition phase, cKO^Htr2b-μglia^_TXFbirth_ male mice learned to choose the correct arm of the Y-maze, containing a food reward, as rapidly as the cWT_TXFbirth_ mice (**Fig. 3o**). By contrast, in the reversal phase of the test, cKO^Htr2b-μglia^_TXFbirth_ male mice failed to learn the new position of the correct arm (**Fig. 3o**), which indicates that early invalidation of 5-HT_2B_ receptors in microglia decreases not only social but also cognitive flexibility.

Sex differences have been reported in many psychiatric disorders, with notably a higher prevalence of neurodevelopmental disorders in males^34^. Also, sex-dependent transcriptional, structural and functional properties of rodent microglia have been described, and this might have potential implications for specific CNS diseases. We thus assessed whether the behavioral phenotypes observed in cKO^Htr2b-μglia^_TXFbirth_ male mice were present also in females. When we assessed their locomotion in a novel environment, adult cKO^Htr2b-μglia^_TXFbirth_ female mice behaved similarly to cWT mice (**Fig. 4a**). Nonetheless, mutant females, similarly to cKO^Htr2b-μglia^_TXFbirth_ males, showed enhanced self-grooming in an unfamiliar cylinder compared to cWT controls (**Fig. 4b**), suggesting an increased propensity to repetitive stereotypic behavior. We also observed a phenotype for juvenile cKO^Htr2b-μglia^_TXFbirth_ female mice in the maternal homing test (**Fig. 4c-d**), similar as the one observed in cKO males: the time spent in the corner containing the tube containing their mother was significantly reduced, and the time spent interacting with the empty tube and in the empty corner were increased, compared to cWT_TXFbirth_ females (**Fig. 4d**). However, we did not observe defects in social behavior in adult females (**Fig. 4e**); this might be due to the different nature of social activity between the two sexes at adulthood, male mice being more sensitive to environmental modifications and more prone to increase their repertoire of social explorations to assess social hierarchy^35^. Finally, we performed reversal learning to assess cognitive flexibility. As for male mice, after a normal acquisition phase, cKO^Htr2b-μglia^_TXFbirth_ females failed to learn the new position of the correct arm in the reversal phase of the test, in comparison with the cWT_TXFbirth_ females (**Fig. 4f**). Of note, since most behaviors required intact olfactory capacities, we tested both genotypes in the olfactory habituation/dishabituation test, and we did not detect any difference in their olfactory skills (males: **Fig. S3a**, females: **Fig. S3b**).

**Fig. 4.**
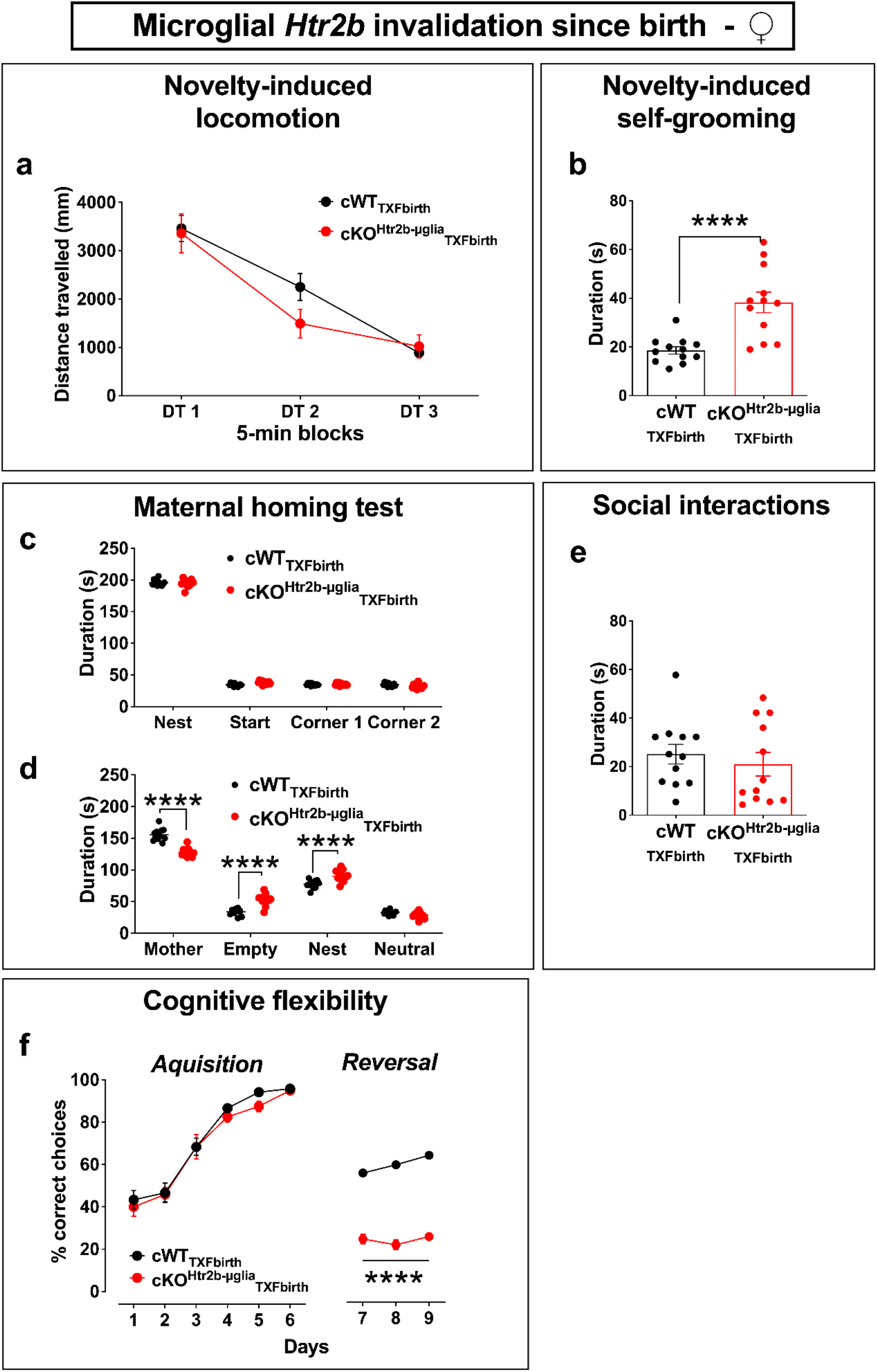
Absence of 5-HT_2B_ receptors in microglia since birth impacts activity of mice in a novel environment, sociability and flexibility in female mice. **a**, Distance travelled in response to a novel environment in 5-min blocks (DT) (two-way RM-ANOVA: significant main effect of DT *P* < 0.0001, *F*_2,44_) = 84.07). **b**, The time spent grooming in response to a novel environment is strongly enhanced in cKO^Htr2b-μglia^_TXFbirth_ adult female mice compared to cWT_TXFbirth_ mice (unpaired *t*-test, p < 0.0001). **c**, Identical time spent in each corner during the initial 5-min habituation period by cWT_TXFbirth_ and cKO^Htr2b-μglia^_TXFbirth_ P25 female mice. **d**, Time spent in each corner during the 5-min of homing test (two-way RM-ANOVA: significant main effects of interaction *P* < 0.0001, *F*_3,66_ = 41.29 and zone *P* < 0.0001, *F*_3,66_ = 933.1); note that the preference for the mother corner was strongly reduced in cKO^Htr2b-μglia^_TXFbirth_ compared to cWT_TXFbirth_ female mice (Sidak multiple comparisons: mother corner, cWT_TXFbirth_ vs. cKO^Htr2b-μglia^_TXFbirth_, p < 0.0001) and, conversely, we observed an increased preference for the empty (Sidak multiple comparisons: empty corner, cWT_TXFbirth_ vs. cKO^Htr2b-μglia^_TXFbirth_, p < 0.0001) and nest corners (Sidak multiple comparisons: nest corner, cWT_TXFbirth_ vs. cKO^Htr2b-μglia^_TXFbirth_, p < 0.0001) in cKO^Htr2b-μglia^_TXFbirth_ mice. **e**, Duration of interaction with a juvenile conspecific in a neutral territory is similar in cWT_TXFbirth_ and cKO^Htr2b-μglia^_TXFbirth_ adult female mice. **f**, During the acquisition phase, all mice reached the criterion to choose the correct arm of the Y-maze; however, cKO^Htr2b-μglia^_TXFbirth_ female mice failed to perform the reversal phase of the test (two-way RM-ANOVA: significant main effects of interaction *P* = 0.0068, *F*_2,44_ = 5.607; phase *P* = 0.0004, *F*_2,44_ = 9.250 and genotype *P* < 0.0001, *F*_1,22_ = 427.3; Sidak multiple comparisons: cWT_TXFbirth_ vs. cKO^Htr2b-μglia^_TXFbirth_, day 7, p < 0.0001; day 8, p < 0.0001; day 9, p < 0.0001). **a-f**: cWT_TXFbirth_, *n* = 12 female mice; cKO^Htr2b-μglia^_TXFbirth_, *n* = 12 female mice. Graphs show mean±s.e.m. and points represent individual animals. ****p < 0.0001.

Overall, early invalidation of *Htr2b* in microglia leads to severe behavioral alteration in adulthood in both sexes, but more pronounced in males, where it is characterized by increased activity (locomotion, self-grooming) in a novel environment, decreased sociability, and deficits in social and cognitive flexibility. These defects could be linked to early microglial and synaptic alterations observed in cKO^Htr2b-μglia^_TXFbirth_ males and females at P15. However, given the importance of 5-HT in behavior, and the expression of 5-HT_2B_ receptor gene in microglia throughout life, we performed additional experiments to disentangle the developmental versus adult (acute) effects of 5-HT signaling in microglia on the phenotype observed.

### 4. Ablation of 5-HT_2B_ receptors in microglia after P30 does not alter spontaneous behaviors

We repeated the same behavioral tests on mice where invalidation of microglial *Htr2b* had been induced at P30, once microglia are mature and synaptogenesis and pruning completed^1^ (cKO^Htr2b-μglia^_TXF-P30_ *vs* CWT_TXF-P30_ mice, both treated with a first dose of tamoxifen between P28 and P30 and a second one 48h later). This protocol was previously shown to be as efficient as the early tamoxifen treatment to induce recombination at the *Htr2b* locus (Fig. S7 in ^11^). Mice were analyzed at least one month after tamoxifen treatment and behavioral experiments were performed separately on males and females (**Fig. 5a**). In this late ablation condition, after checking normal olfactory skills (males: **Fig. S3c**; females: **Fig. S3d**), we did not detect any deficit in locomotion or selfgrooming in a new environment (males: **Fig. 5b-c**; females: **Fig. S4a-b**), nor in sociability (males: **Fig. 5d**; females: **Fig. S4c**), or in social (males: **Fig. 5e-f**) or cognitive flexibility (males: **Fig. 5g**; females: **Fig. S4d**). This is consistent with the idea that these behavioral domains rely on the establishment of a proper brain connectivity during early postnatal development^36^. Besides, these results indicate that the behavioral defects observed when microglial *Htr2b* is invalidated since birth are directly linked to an impairment of serotonergic signaling to microglia during a postnatal window between birth and P30.

**Fig. 5.**
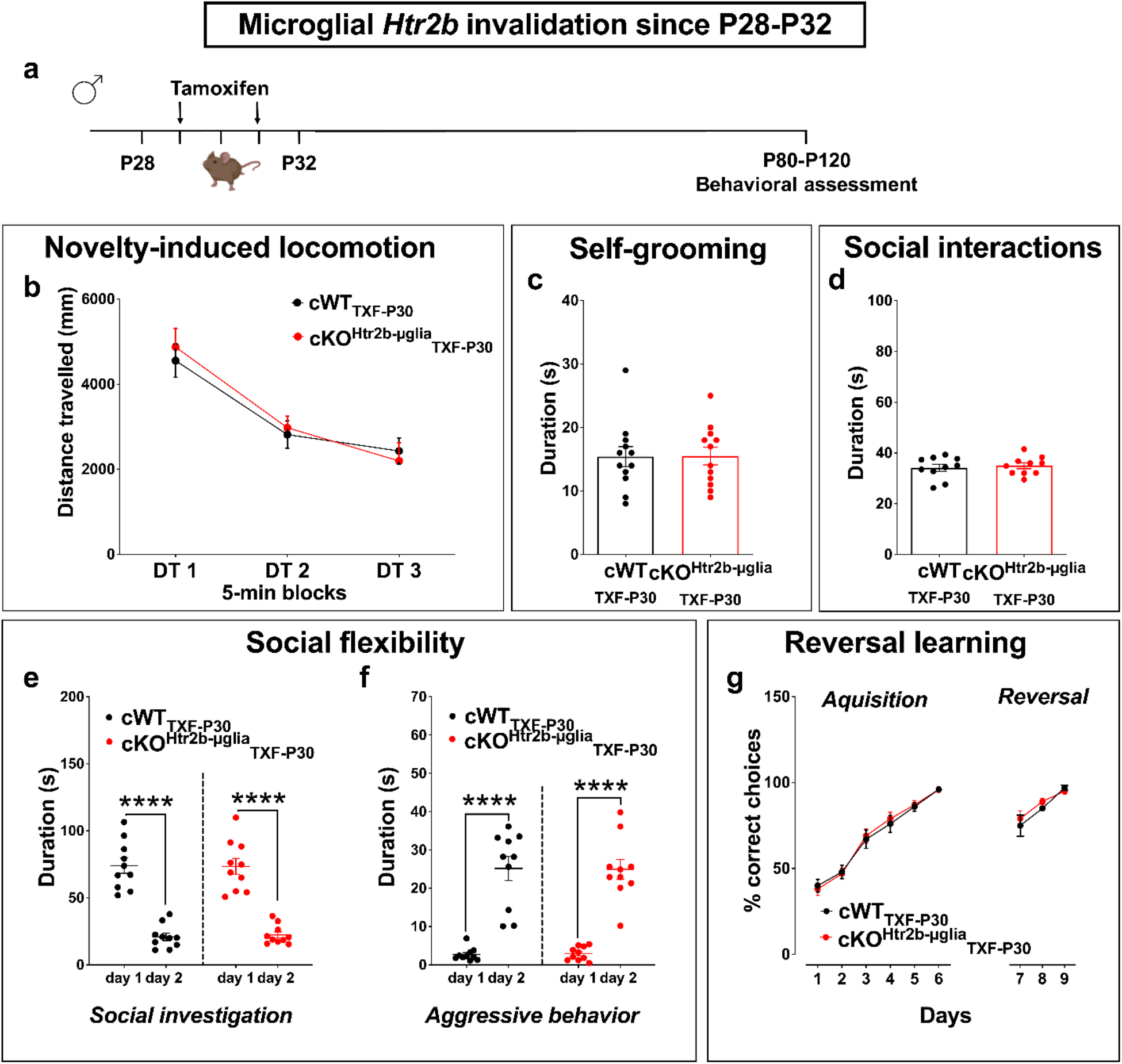
Ablation of 5-HT_2B_ receptors in microglia after P30 does not alter spontaneous behaviors in adult male mice. **a,** Timeline of tamoxifen treatment for late microglial *Htr2b* invalidation (P28-32), and the behavioral tests performed in adult male mice. **b**, Similar distance travelled in response to a novel environment in 5-min blocks (DT) (two-way RM-ANOVA: significant main effect of DT *P* < 0.0001, *F*_2,36_ = 70.78). **c**, Similar time spent self-grooming in response to a novel environment in CWT_TXF-P30_ and cKO^Htr2b-μglia^_TXF-P30_ adult male mice. **d**, Similar time spent interacting with a juvenile conspecific in a neutral territory in cWT_TXF-P30_ and cKO^Htr2b-μglia^_TXF-P30_ adult male mice. **e-f**, Spontaneous social behavior in the home cage of adult male mice housed with a familiar cage mate, during two consecutive days, using the change of litter bedding on day 2 as an environmental challenge. **e**, Duration of social investigation (two-way RM-ANOVA: significant main effect of day *P* < 0.0001, *F*_1,18_ = 203.7); note that both CWT_TXF-P30_ and cKO^Htr2b-μglia^_TXF-P30_ adult male mice modified their behavior in agreement to the change in social hierarchy induced by environmental challenge (Sidak multiple comparisons: CWT_TXF-P30_, day 1 vs. day 2, p < 0.0001; cKO^Htr2b-μglia^_TXF-P30_, day 1 vs. day 2, p < 0.0001). **f**, Duration of aggressive behavior (two-way RM-ANOVA: significant main effect of day *P* < 0.0001, *F*_1,18_ = 145.8); note that both CWT_TXF-P30_ and cKO^Htr2b-μglia^_TXF-P30_ adult male showed increased aggressive behavior after the change of the bedding (Sidak multiple comparisons: CWT_TXF-P30_, day 1 vs. day 2, p < 0.0001; cKO^Htr2b-μglia^_TXF-P30_, day 1 vs. day 2, p < 0.0001). **g**, Y-maze reversal learning shows no difference in cognitive flexibility task in cWTTXF-P30 and cKO^Htr2b-μglia^TXF-P30 adult male mice. **b, d-g**: cWT_TXFbirth_, *n* = 10 male mice; cKO^Htr2b-μglia^_TXFbirth_, *n* = 10 male mice. **c**: cWT_TXFbirth_, *n* = 12 male mice; cKO^Htr2b-μglia^_TXFbirth_, *n* =12 male mice. Graphs show mean±s.e.m. and points represent individual animals. ****p < 0.0001.

## Discussion

In this work, our aim was to investigate if disturbed sensing of 5-HT by microglia affects the microglial and neuronal maturation, and can be implicated in behavioral alterations observed in neurodevelopmental psychiatric disorders. Our findings demonstrate that interrupting the control of microglia by 5-HT perturbs microglial functions during postnatal development, affects synaptic and axonal refinement throughout the brain at P15-P18, and induces long-lasting behavioral phenotypes in adulthood such as repetitive behavior in a new environment, deficits in sociability and a lack of behavioral flexibility. Importantly, these behavioral alterations, which are reminiscent of core symptoms of neurodevelopmental disorders are not induced by the ablation of microglial 5-HT_2B_ receptors after P30. This work thus provide evidence that behavioral features (including response to novelty, sociability and flexibility) require the developmental control of 5-HT on microglia.

### Microglial 5-HT_2B_ receptor is required in the neonatal period for proper microglial functions and brain wiring

Our data demonstrate that early (i.e. since birth) invalidation of microglial *Htr2b* triggers higher density of dendritic spines and enhanced excitatory neurotransmission in the hippocampus at P15. Synaptic refinement occurs early in postnatal development and relies, among other mechanisms, on active contacts between microglia and dendritic shafts, promoting spine formation^21,30^, and on microglial phagocytosis of presynaptic structures^13,21–23^. Intriguingly, if microglia cannot detect early 5-HT signal through 5-HT_2B_ receptors, they display reduced CD68^+^ phagolysosomal volume at P15, suggesting reduced phagocytic capacity in physiological condition. In addition to the phenotype observed in hippocampal and cortical neurons, presynaptic contralateral and ipsilateral inputs in the dLGN are still intermingled in cKO^Htr2b-μglia^_TXFbirth_ mice at P18, resembling earlier stages of postnatal development. In line with their decreased CD68 content, microglia invalidated for *Htr2b* might be less prone to eliminate presynaptic inputs in the dLGN. Additionally, we discovered that a reduced fraction of hippocampal spines is found in close proximity of microglial processes in cKO^Htr2b-μglia^_TXFbirth_ mice, suggesting that the absence of 5-HT signal in microglia alters their ability to interact with surrounding neurons. Given that microglia provide a negative feedback control of neuronal activity^37^, reduced proximity between microglia and neurons could directly contribute to the enhanced excitatory neurotransmission observed in cKO^Htr2b-μglia^_TXFbirth_ mice.

### Microglial 5-HT_2B_ receptor supports serotonin-dependent refinement of brain wiring during postnatal development

It was previously reported that segregation of retinal projection into eye-specific territories in the dLGN does not occur in monoamine oxidase A knock-out mice (MAOA-KO), which have elevated brain levels of 5-HT, and that these defects can be reversed by inhibiting 5-HT synthesis from P0 to P15. The serotonin transporter (SERT), which is transiently expressed in the dLGN into the ipsilateral retinal projection fields, has been proposed to confer specific neurotransmission properties on a subset of retinal ganglion cells that are important for proper segregation of retinal projections^14^. This was reinforced by the complementary evidence that eye-specific segregation in the dLGN and topographic refinement of ipsilateral axons in the dLGN are impaired in mice lacking vesicular release, in RIM1/2 conditional knock-out mice in SERT-positive neurons^38^. We previously reported that full 5-HT_2B_ receptor deficiency induces anatomical alterations of the ipsilateral projecting area of retinal axons into the dLGN at P30, validating the contribution of this receptor into 5-HT-dependent retinal projection segregation^9^. Here, we demonstrate that presynaptic contralateral and ipsilateral inputs in the dLGN are still intermingled in cKO^Htr2b-μglia^_TXFbirth_ mice at P17-18, revealing that part of the 5-HT contribution to this segregation is to control microglia.

### Microglial 5-HT_2B_ receptor is required during the neonatal period, but not later, for adult social behavior and cognitive flexibility

We previously reported defects in social interactions in *Htr2b^-/-^* male mice^16^. Hereby, we observed that, if microglial 5-HT_2B_ receptors are absent since birth, both male and female mice later display a plethora of behavioral abnormalities resembling key symptoms of neurodevelopmental psychiatric disorders, notably abnormal response to novelty, and decreased sociability and adaptability. Intriguingly, these behavioral alterations were not induced by the absence of these receptors after P30. The evidence that invalidation of 5-HT_2B_ receptors in microglia specifically since birth leads to such dramatic and long-lasting consequences, suggests that 5-HT acts as a global modulator of microglia during the critical time-window of postnatal development before P30, which is required for the establishment of specific behavioral domains.

### Sex-dependent effects of the invalidation of microglial 5-HT_2B_ receptors

Microglia exhibit sexually dimorphic transcriptomic and morphological differences along postnatal development and adulthood^28,39–41^. Since neurodevelopmental and psychiatric disorders show a marked sexual dimorphism, it is plausible that sex-dependent microglial functions contribute to the different prevalence of these diseases among males and females. As well, a PET scan study demonstrated that women’s brains show lower 5-HT activity^42^, possibly reflecting 5-HT contribution to sex differences in psychiatric disorders. We therefore paid attention to possible sex-dependent effects of microglial 5-HT_2B_ receptors invalidation, demonstrating that early invalidation of microglial *Htr2b* induces alteration of the microglial morphology (reduced complexity of the ramifications) in male mice only. However, apart from this morphological readout, we did not observe sex differences in the effects of early invalidation of microglial 5-HT_2B_ receptors on microglial CD68 content, nor synaptic refinement and behavior, which highlights the universal importance of 5-HT sensing by microglia during postnatal development.

### Mechanisms linking activation of microglia through 5-HT_2B_ receptors and downstream effects

A major, but still open, question concerns the intracellular signaling effectors of 5-HT in microglia. Both 5-HT and ATP/ADP, activating P2Y12 receptors, induce directional motility of microglial processes^9,12,43^. Moreover, pharmacologic or genetic P2Y12 receptors disruption in adolescent mice induces a reduction in microglial ramification^44^, similar to what we observed in males mice invalidated for *Htr2b*. It is thus possible that G_i_-coupled P2Y12- and G_q_-coupled 5-HT_2B_ receptor-mediated signals are interconnected, either physically, as GPCR heterodimers^45^, or functionally. This latter possibility would remind of a mechanism existing in platelets, where aggregation is induced either by ADP^46^ or 5-HT^47^ through activation of P2Y12 or 5-HT_2A_ receptors-which is structurally very close to 5-HT_2B_ receptors-, respectively, and inhibited by P2Y12 or 5-HT_2A_ receptor antagonists^48,49^. Our results also raise the question of how the stimulation of microglia by 5-HT impacts on synapses and neurons. Our findings suggest that during development, reduced microglial phagocytic capacity and proximity with spines in the absence of 5-HT signaling in microglia is responsible for the altered refinement and thus the behavioral impairments. However, additional mechanisms might be involved, such as decreased somatic contacts or altered secretion of factors affecting spine plasticity.

In summary, our data strongly support that 5-HT acts as an upstream global modulator of microglial functions during early postnatal development, and that a lack of serotonergic control of microglia affects brain connectivity refinement and thereby a variety of behaviors that are altered in neurodevelopmental disorders. Given the number of exogenous factors known to change 5-HT levels during postnatal development, such as physical abuse, maternal separation and maternal inflammation, stress, antidepressants, and their correlation to neurodevelopmental disorders, our findings further emphasize the need of optimal brain levels of 5-HT for proper brain development and function throughout life. Besides, as the serotonergic system is targeted by an array of pharmacological compounds, this newly identified pathway of microglia regulation might open new therapeutic avenues for early intervention in neurodevelopmental disorders.

## Materials and methods

### Animals

Animal experiments were carried out according to the national guidelines on animal experimentation and were approved by the Ethical Committee for Animal Experiments of the Sorbonne University, Charles Darwin C2EA - 05 (authorization N°APAFIS#28230-2020111619109364 v4, and 01170.02). All efforts were made to minimize animal suffering. Results are described in accordance with the ARRIVE guidelines for reports in animal research^50^. To obtain targeted invalidation of *Htr2b* gene in microglia, we crossed the *Cx3cr1*^creERT2^ line (MGI Cat# 5528845, RRID:MGI:5528845), which express a tamoxifen-activable cre recombinase under the control of the endogenous *Cx3cr1* promoter, and the *Htr2b^flox^* line (*Htr2b^tm2Lum^/Htr2b^tm2Lum^, IMSR Cat# EM:05939, RRID:IMSR_EM:05939*), where loxP sites are inserted around the first coding exon of *Htr2b* gene, allowing invalidation of the gene when the recombinase is activated by tamoxifen as previously described^11,51,52^. All lines were backcrossed on 129S2 background for more than 10 generations. *Cx3cr1^creERT2/+^;Htr2b^fl/fl^* mice were used to obtain the conditional invalidation (see below the protocol to activate the creERT2 recombinase with tamoxifen). *Cx3cr1^creERT2/+^;Htr2b^+/+^* mice, similarly treated with tamoxifen, were used as control mice. P15, P25 or P80-120 (adult) male and female animals were used for the experiments (see the experimental design below and **Table 1**). All mice were group-housed (2–5 mice per cage) in a 12/12 h light/dark cycle with free access to food and water. Temperature and relative humidity were maintained at 21±1°C and between 30% and 60%, respectively.

**Table 1.** Tamoxifen administration and experimental design.

| **Time of invalidation of *Htr2b* in microglia** | **Age at the time of the analysis** | **Sex** | **Analyses performed** |
| --- | --- | --- | --- |
| "Early" invalidation (tamoxifen P1-P5), i.e. "TXFbirth" subscript | P15 | Males and females | Immunohistochemistry to study microglia, DiI/P2Y12 receptor labeling, Golgi staining, patch clamp recordings (*ex vivo*) |
|  | P15-P18 | Males and females | Anterograde labeling (P15) and analysis (P18) of retinogeniculate projections (*ex vivo*) |
|  | P25 (maternal homing test), P80-P120 | Males and females | Behavioral assessment |
| "Late" invalidation (tamoxifen P28-P32), i.e. "TXF-P30" subscript | P80-P120 | Males and females | Behavioral assessment |

### Tamoxifen administration

To study the consequences of invalidation of microglial *Htr2b* at different time points, we used the following experimental approaches. First, to invalidate the *Htr2b* gene since birth (“early” invalidation), we performed intragastric administrations of 50 μg of tamoxifen (Sigma-Aldrich, T5648) per day for 3 days between P1 and P5 as previously described^53^. In a second approach, to invalidate the *Htr2b* gene since P30 (“late” invalidation), animals were fed twice via oral gavage with a 20 mg/ml tamoxifen solution between P28 and P32, at 48 h interval, as previously described^51^. At P28-32, the mice weighed ~ 15–25 g and received 200–400 μl, corresponding to 0.2 mg.g^-1^ body weight. Of note, we previously validated the efficacy of our protocols to activate creERT2 recombinase by determining by qPCR the level of *Htr2b* recombination in the microglial fraction of mice that received early or late tamoxifen administration, demonstrating a similar decrease (>85%, Figure S7C in ^11^). Moreover, we waited at least 1 month after tamoxifen treatment to allow the full renewal of peripheral Cx3cr1^+^ cells^51,54–57^.

### Immunohistochemistry

Mice were deeply sedated (pentobarbital, 140 mg.kg^-1^ i.p.) and transcardially perfused with 4% formaldehyde in PBS. Brains were harvested and immersed in the same fixative for 24 h at 4°C. Afterwards, they were transferred to a PBS solution with 0.01% sodium azide until being cut. Coronal slices were cut using a vibratome (Leica). Four to six sections per mouse were selected around 1.82 mm caudal to bregma following the mouse brain atlas of Franklin and Paxinos^58^. Brain slices (50 μm for microglial analysis and 40 μm for c-fos^+^ cells density) were washed one time in 0.1 M PB, then incubated in permeabilizing and blocking solution (0.25% gelatin and 0.1% triton in PB) for 30 min. Primary antibodies, rabbit anti-Iba1 (0.0013 μg.μL^-1^, Wako 019-19741), rat anti-CD68 (0.002 μg.μL^-1^, AbD Serotec MCA1957), and rat anti-P2Y12 receptor (0.5 μg.μL-1, Biolegend 848002) were diluted in PGT buffer (0.125% gelatin and 0.1% triton in PB). Samples were incubated for 24 h at 4°C with gentle agitation. After rinsing three times with PGT 0.125% buffer, secondary fluorochrome-conjugated antibodies were added to PGT buffer for 3h at room temperature. To quantify Iba1^+^ cells density, we used a donkey anti-rabbit Cy3 (0.002 μg.μL^-1^, Jackson Laboratories 711-165-152); to colocalize Iba1/CD68 and to assess Iba1^+^ cells morphology, we used donkey anti-rabbit 488 (0.002 μg.μL^-1^, Invitrogen A21206) and donkey anti-rat Cy3 (0.002 μg.μL^-1^, Millipore AP189C). After rinsing three times with PGT 0.125% buffer and once with PB, sections were mounted in Mowiol containing DAPI (0.005 μg.μL^-1^, Sigma).

### Image acquisition and analysis

For quantification of Iba1^+^ cell density, images were acquired with a Leica DM6000 epifluorescence microscope equipped with a CCD camera (CoolSNAP EZ, Teledyne Photometrics). Density of Iba1^+^ cells was quantified on 20x images collected in several regions (identified anatomically), including CA1, CA3 and dentate gyrus (DG) of the dorsal hippocampus, retrosplenial cortex (Ctx) and dLGN. For each region, one image per hemisphere was taken from six separate brain sections to obtain an average value for that mouse. Images were smooth processed and binarized, and Iba1^+^ cells were identified and counted using the Analyze particle plugin for ImageJ, setting the minimum detectable size at 40 μm^2^.

For 3D reconstruction of entire microglial cells and analysis of microglial lysosome content, confocal fluorescent images were acquired on an upright confocal microscope (Leica TCS SP5), using a 40x/1.25-0.75 oil (11506253) objective with 2X electronic magnification (188.92 nm pixel size) and lasers set at 488 and 561 nm for excitation of FITC and Cy3, respectively. Stacks were acquired using a 0.5 μm z-interval. Image stacks were imported into ImageJ for processing as previously described^59,60^. Briefly, to edit the channel window containing Iba1^+^ cells, an Unsharp Mask filter was used to further increase contrast using a pixel radius of 3 and mask weight of 0.6, a despeckle step was performed to remove salt-and-pepper noise and outlier pixels were removed with the Remove Outliers function (pixel radius 2; threshold 50). Image stacks were then imported into Imaris Software (Bitplane). 3D reconstructions of individual microglia were carried out using the filament tracer module and measures of the process length, terminal points, as well as 3D-Sholl analysis were performed to assess microglia morphology. For analysis of intracellular CD68 distribution, surface module was used to reconstruct the volume of microglia and CD68-labeled lysosomes, and then the percent of Iba^+^ cell occupied by CD68 was assessed as previously described^59^. The experimenter carrying out the acquisition and analysis of images was blind to the mice genotype.

### DiI labeling of dendritic spines and analysis of microglial proximity

Deeply sedated mice (pentobarbital, 140 mg.kg^-1^ i.p.) were transcardially perfused with 2% formaldehyde in PBS. Whole brains were removed, immersed in the same fixative for 1 h, transferred to a solution consisting of PBS with 0.01% sodium azide until being cut, and sectioned at 100 μm thickness on a vibratome (Leica). Two sections per mouse were selected around 1.82 mm caudal to bregma following the mouse brain atlas of Franklin and Paxinos^58^. DiI labeling of dendritic spine and spine analysis was performed as previously described^61^. Briefly, DiI beads (1,10-dioctadecyl-3,3,30,30-tetramethylindocarbocyanine perchlorate; Molecular Probes, Eugene, OR) were delivered to the slices using a helium gas pressure (100–150 psi) through the gene gun device (BioRad). A 3 μm pore size filter (Isopore polycarbonate, Millipore) was inserted between the gene gun and the slice to select single beads. Slices were and kept in PBS at room temperature for at least 2 h to allow DiI diffusion in neurons, then incubated in permeabilizing solution (0.1% triton in phosphate buffer, PB) for 5 min, followed by blocking solution (0.25% gelatin and 0.01% triton in PB) for 45 min. Rat anti-P2Y12 receptor (0.5 μg.μL^-^1, Biolegend 848002) was diluted in a solution of 0.125% gelatin and 0.01% triton in PB and samples were incubated for 72 h at 4°C with gentle agitation. After rinsing three times with 0.125% gelatin and 0.01% triton in PB, donkey anti-rat 488 (0.0013 μg.μL^-^1, Invitrogen A21208) was added to the buffer for 3h at room temperature. Slices were mounted in Mowiol containing DAPI (0.005 μg.μL^-1^, Sigma). Image stacks of apical dendritic segments in CA1 were taken using a confocal laser scanning microscope (SP5, Leica) equipped with a 63x/1.4 NA objective (oil immersion, Leica) with a pinhole aperture set to 1 Airy unit, pixel size of 60 nm and z-step of 200 nm. The excitation wavelength and emission range were 488 and 500-550 nm for Alexa 488 (i.e. P2Y12 receptor) and 561 and 570-620 for DiI. Deconvolution with experimental point spread function from fluorescent beads using a maximum likelihood estimation algorithm was performed with Huygens software (Scientific Volume Imaging). One hundred fifty iterations were applied in classical mode, background intensity was averaged from the voxels with lowest intensity, and signal to noise ratio values were set to a value of 20. For each mouse, 10 dendritic segments of 20–50 μm in length were analyzed. Dendritic spines were analyzed with Neuronstudio software^62^ (version 9.92; http://research.mssm.edu/cnic/tools.html). Images from the P2Y12 receptor channel were preprocessed in order to enhance the contrast, using an Unsharp Mask filter using a pixel radius of 3 and mask weight of 0.6, a despeckle step was performed to remove salt-and-pepper noise and outlier pixels were removed with the Remove Outliers function (pixel radius 2; threshold 50). To analyze the interaction between microglial processes and dendritic spines, we measured the percentage of dendritic spines having microglial processes located within 0.3 μm surrounding the spine head using Fiji-ImageJ software. To assess the presence of microglial processes, for each spine, its 3D coordinates and head diameter measured with Neuronstudio were used to draw a sphere on the image using the DrawShape function of the plugin 3DImageSuite. A second sphere centered on the spine coordinates with a diameter extended by 0.3 μm was then drawn. The volume space in between the two spheres was converted into a Region of Interest. The mean intensity inside the 3D ring volume was assessed in the channel corresponding to Alexa 488 (i.e. P2Y12 receptor), and compared to that of the background. If the mean intensity was more than ten times than that of the background, it was considered that the spine had microglial processes in proximity.

### Golgi staining and spine analysis

Fresh brains were processed following the Golgi-Cox method as described before^63,64^. Briefly, mice were deeply sedated (pentobarbital, 140 mg.kg^-1^ i.p.) and brains were harvested, briefly rinsed in NaCl, and incubated in the dark in filtered dye solution (10 g.L^-1^ K_2_Cr_2_O_7_, 10 g.L^-1^ HgCl_2_ and 8 g.L^-1^ K_2_CrO_4_) for 15-16 days. Impregnated brains were then washed 3 × 2 min in distilled water (dH_2_O) and transferred to a tissue-protectant solution (30% sucrose in dH_2_O) for 24 h. Afterwards, brains were rinsed 3 x 2 min in dH_2_O and incubated 30 min in 90% EtOH (v/v). Two hundred μm-thick coronal sections were cut in 70% EtOH on a vibratome (Leica) and washed in dH_2_O for 5 min. Next, sections containing the dorsal hippocampus (6 sections per mouse) or medial prefrontal cortex (6 sections per mouse) were reduced in 16% ammonia solution for 1 h, washed in dH_2_O for 2 min and fixated in 10 g.L^-1^ Na_2_S_2_O_3_ for 7 min. After a final wash in dH_2_O, sections were mounted on superfrost slides and allowed to dry at room temperature for 1-2 h. The dehydration steps proceeded as following: 3 min in 50% EtOH, 3 min in 70% EtOH, 3 min in 80% EtOH and 3 min in 100% EtOH, 2 × 5 min in a 2:1 isopropanol:EtOH mixture, 5 min in pure isopropanol and 2 × 5 min in xylol. Neurons were first identified with a light microscope (Leica 6000) under low magnification (×20). At least three neurons per mouse were selected according to the following criteria: (1) consistent and dark impregnation along the entire extent of all the dendrites; (2) presence of fourth-order branches for both apical and basal dendrites, (3) presence of untruncated dendrites, (4) soma entirely within the thickness of the section, and (5) relative isolation from neighboring impregnated neurons. Bright-field images of z-stacks of apical dendrites from hippocampal CA1 and L2/L3 pyramidal neurons were then captured with a × 100 oil objective every 0.2 μm using a CoolSNAP EZ CCD Camera (Teledyne Photometrics). Given that protrusions density reached a plateau 45 μm away from the soma, we selected dendritic segments at least 50 μm away from the cell body. Protrusions were counted on secondary branches of apical dendrites. Only protrusions with a clear connection of the head of the protrusion to the shaft of the dendrite were counted. Protrusion density, length and width were analyzed using the Reconstruct software as described by Risher and colleagues^31^. Protrusions were classified based on their morphology into six classes using the following exclusive criteria^31^: filopodia, when the length value >2 μm; long thin spines, when the length value >1 μm and ≤2 μm; thin spines, when the length-to-width ratio (LWR) value >1; mushroom spines, when the width value >0.6 μm; stubby spines, when the LWR value ≤1; branched spines, when spines showed more than one head (entered manually by the experimenter). All image analyses were done blind.

### Patch clamp recordings

Two hundred fifty μm-thick transverse hippocampal slices were prepared from brains of P15 mice. Mice were deeply sedated (pentobarbital, 140 mg.kg^-1^ i.p.) and the brain was quickly removed from the skull, then sectioned with a vibroslicer (HM 650 V, Microm) in ice-cold artificial cerebrospinal fluid (ACSF) containing the following (in mmol.L^-1^): 125 NaCl, 2.5 KCl, 25 glucose, 25 NaHCO_3_, 1.25 NaH2PO_4_, 2 CaCl_2_, and 1 MgCl_2_, continuously bubbled with 95% O2-5% CO_2_. Slices were incubated in ACSF at 32°C for 20 min and then at room temperature (20-25°C). For patch-clamp recordings, slices were transferred to the recording chamber where they were continuously superfused with ACSF (30-32°C) buffered by continuous bubbling with 95% O_2_-5% CO_2_. Patch-clamp pipettes (4-6 Mohm resistance) were prepared from borosilicate glass (BF150-86-10; Harvard Apparatus) using a DMZ pipette puller (Zeitz). CA1 pyramidal neurons were visually identified using an upright microscope (Olympus BX51WI). Neurons were voltage-clamped using an EPC10 amplifier (HEKA Elektronik) and the following intracellular solution (in mmol.L^-1^): 120 Cs-methane sulfonate, 10 CsCl, 10 Hepes, 10 Phosphocreatine, 0.2 EGTA, 8 NaCl, 2 ATP-Na_2_, 3 QX 314 (pH 7.25, adjusted with CsOH). Miniature excitatory postsynaptic currents (mEPSCs) were recorded in CA1 pyramidal neurons at a holding potential of −65 mV in the presence of Tetrodotoxin (TTX, 0.5 μM, Hello Bio) and SR95531 hydrobromide (Gabazine, 10 μM, Hello Bio). At least 200 s of recording were analyzed for each cell. Series resistance was left uncompensated. Input resistance was monitored by voltage steps of −10 mV before and after mEPSC recordings. Experiments were discarded if the series resistance changed by more than 20% during the recording. Data acquisition was performed using Patchmaster software (Heka Elektronik). The junction potential (−14.9 mV) was uncorrected. Signals were low pass-filtered at 4 kHz before sampling at 20 kHz. Recordings were filtered offline at 2 kHz using Clampfit (Molecular Devices) and analyzed using MiniAnalysis (Synaptosoft). During patch-clamp recordings and analysis, the investigator was blind to the mouse genotype.

### Anterograde labeling of retinogeniculate projections

P15-16 mice were anesthetized with ketamine-xylazine (100 and 10 mg.kg^-1^, respectively, in 0.9% saline). Eyes were intravitreally injected using a glass micropipette with 1-5 μl of 0.2% cholera toxin subunit B (CTB) conjugated to Alexa Fluor 488 or 555 (Invitrogen) diluted in 1% DMSO. After 48 h, mice were deeply sedated (pentobarbital, 140 mg.kg^-1^ i.p.) and perfused transcardially with 4% paraformaldehyde in 0.1 M phosphate buffered saline (PBS). Brains were postfixed overnight with the same fixative and sectioned with a vibratome (60 μm). All sections containing the dorsal lateral geniculate nucleus of the thalamus (dLGN) were selected and directly mounted in Mowiol containing DAPI (0.005 μg.μL^-1^, Sigma). 10X epifluorescent images were obtained using a CCD camera (CoolSNAP EZ, Teledyne Photometrics) attached to an upright Leica 6000 microscope.

MetaMorph software (Molecular Devices) was used for quantitative analyses. All image analyses were done blindly to the mice genotypes. Quantification of eye-specific segregation was performed on images of Alexa Fluor 488-labeled contralateral/Alexa Fluor 555-labeled ipsilateral projections, as previously described by Rebsam and colleagues^65^. Briefly, variable thresholds were used for the contralateral projection and a fixed threshold for the ipsilateral projection. After outlining the boundary of the dLGN, we measured the pixel overlap between ipsilateral and contralateral projections at every 10th threshold value for the contralateral image. The proportion of dLGN occupied by ipsilateral axons was measured as a ratio of ipsilateral pixels to the total number of pixels in the dLGN region. Additionally, we quantified the areas occupied by ipsilateral and contralateral retinogeniculate projections and the extent of ipsilateral retinogeniculate projections along the outer-inner axis (O-I axis), perpendicular to the surface of the dLGN, and the dorsomedial-ventrolateral axis (DM-VL axis), parallel to the surface of the dLGN, as previously described by Hayakawa and Kawasaki^66^. The extent of the ipsilateral projection along the O-I and DM-VM axes was measured by tracing a line between the two points delineating the maximal extent of the ipsilateral signal along the chosen axis. The extent of the ipsilateral projection along the dLGN axis is calculated by dividing the length of the ipsilateral projection by the dLGN length and is expressed as a percentage of dLGN length. For quantifying the areas occupied by ipsilateral retinogeniculate projections, thresholds (Ti) were determined as follows: Ti = Mi x 0.165, where Mi was the maximum signal intensity in five consecutive sections, the third of which was the section containing the largest dLGN area. The size of the area whose signal intensities were more than Ti was used as the size of ipsilateral retinogeniculate projections. For quantifying the areas occupied by contralateral retinogeniculate projections, thresholds (Tc) were determined as follows: Tc = Mc x 0.5, where Mc was the average signal intensity of contralateral retinogeniculate projections. The areas whose signal intensities were less than Tc consisted of two kinds of regions: the “gap,” which is the CTB-negative area in the central dLGN, and the regions that are connected with the contours of the dLGN. In order to measure the size of the gap, the latter was excluded.

### Behavioral assessment

Experiments were carried out in the light phase of the light/dark cycle (between 8:00 a.m. and 8:00 p.m.). For each behavioral test, mice were acclimatized to the testing room for at least 1 hr. Equipment was carefully swabbed with 35% ethanol between each mouse to eliminate odors. Behavioral tests were video-recorded in real-time and analyzed off-line by an experimenter blinded to the experimental groups using the free event recorder JWatcher software (Blumstein & Daniel http://www.jwatcher.ucla.edu/).

#### Novelty-induced locomotion

Adult mice were individually placed in a cage with the same size of the home-cage containing a thin layer of fresh bedding, and allowed to freely explore for 15 min. Total distance moved and mean velocity were scored during 3 5-min blocks.

#### Novelty-induced self-grooming

Adult mice were individually placed in a Plexiglas cylindrical arena (20 cm in diameter), and the total time spent performing face, body and/or tail grooming was scored for 5 min as previously described^67^.

#### Homing test

P25 mice were subjected to a maternal homing test as previously described^68^. This test exploits the tendency of mice to maintain body contact with the mother and their siblings and tests olfactory, visual and motor capacities. Mice were separated from the mother for at least 30 min before testing. During habituation, individual juveniles were transferred to a Plexiglas arena (23 × 23 cm, walls 13 cm high) containing fresh bedding with a small amount of soiled nest (not including fecal boli) sprinkled into the opposite corner and allowed to explore for 5 min. Total time spent in the starting, nest and neutral corners was measured. Subsequently, the mice were briefly removed to a holding cage while two wire-mesh tubes were placed into the neutral corners, one empty and one containing the animal’s mother. Time spent in the four corners as well sniffing the tubes (empty vs. mother) was scored.

#### Social interaction

Each experimental mouse was individually transferred to a novel cage. After 15 min of habituation, an unfamiliar juvenile mouse (7 weeks old) of the same sex and strain, used as a social stimulus, was introduced in the cage for 10 min. The following behavioral patterns were used to define social investigation: anogenital sniff (sniffing the anogenital area of the partner); nose sniff (sniffing the head and the snout region of the partner); body sniff (sniffing any other area of the body of the partner); allogrooming (grooming the partner). The duration of social investigation was measured.

#### Social flexibility

Social and aggressive behavior in response to a challenge in social hierarchy was recorded according to the paradigm previously described^69,70^. Briefly, mice housed two by two of the same genotype were observed in their home cage, for a single 30 min session on two consecutive days. During the first day of observation, no manipulation was performed. On the second day, the cage sawdust was changed with clean sawdust just before the session of observation. This procedure is known to elicit aggressive behavior and to challenge social hierarchy. The behavioral elements scored for duration were related to social investigation (sniffing and grooming the partner in all body regions indicative of affiliative behavior) or aggressive behavior (active fighting episode of attacks and aggressive grooming, as well tail rattling and digging behaviors aimed to assess dominant status). Since the data collected was dependent, only one mouse, randomly selected from each cage, has been considered in the analysis. In order to be able to discriminate the two animals during data collection, seven days before the first day of observation, the cages were cleaned and one of the two experimental subjects was marked with a blue, scentless and nontoxic felt pen.

#### Reversal learning

To test memory formation and cognitive flexibility, mice were subjected to a Y-maze task. The apparatus consisted of a transparent, Plexiglas Y maze, composed of 3 test arms forming the Y. The entire procedure took 10 consecutive days: 1 day of habituation, 6 days of acquisition and 3 days of reversal learning. One of the arms was randomly selected as starting arm and, during habituation, mice were individually located in the maze, free to explore for 30 min. In the acquisition phase, one of the other two was baited with a food reward pellet (Noyes sucrose pellet, 20 mg, Research Diets, Inc., New Brunswick, NJ). The side of the rewarded arm was balanced across mice in order to avoid any side preference. Mice were trained to find the food reward in 10 consecutive trials for 6 days. If the mouse made the correct choice, it was given time to consume the sugar pellet, and then guided back into the start arm for the next trial. Incorrect choices were not rewarded or punished. If the mouse did not make any choice in 3 min, it went back to the home cage and the trial was considered as incorrect. For each successive trial, the reward was always placed in the same arm. Number of errors in arm selection, and number of days to criterion were recorded by an observer, blind to the genotypes. The % of correct choice of one session (i.e., day) is the ratio of the number of correct choices divided by the number of trials (=10). The criterion for task acquisition, determined for each mouse, was 80% correct responses on three consecutive days. Each mouse that met criterion for acquisition was then further tested using a reversal procedure, in which the reward location was switched to the arm opposite to its previous location for each mouse. Ten trials per day were then performed for reversal learning, using the same methods and criterion as described above. Subjects were food-restricted, maintaining them at 80-85% of their individual body weight calculated under ad libitum feeding conditions, starting 1 week before the beginning of the experiment and throughout the entire procedure.

#### Olfactory habituation/dishabituation test

The ability to discriminate non-social and social odors was measured using modifications of the olfactory habituation/dishabituation task, as previously described^73^. Subjects were individually tested for time spent sniffing cotton tipped swabs suspended from the cage lid. The olfactory cues were designed to measure familiar and unfamiliar odors, with and without social valence. Sequences of three identical swabs assayed habituation to the same odor, resulting in a progressive decrease in olfactory investigation. Switching to a different odor on the swab assayed dishabituation, i.e. recognition that an odor is new. Five different odors were presented in three consecutive trials of 2 min per trial, with an inter-trial interval of 1 min. The order of swaps was: water (neutral odor), two non-social odors (almond and banana) and two social scents (social 1, unfamiliar mouse of the same sex and different strain, and social 2, unfamiliar mouse of the same strain and opposite sex). In each session trials 50 μL H_2_O or 50 μL of a 1:100 dilution of the odor solutions in H_2_O were pipetted onto a new cotton applicator directly before the test and sealed in a closed tube until the presentation. Unfamiliar social scents were prepared on the test day by swapping the bottom of a cage with a mouse from a different strain (i.e. C57BL/6J) or opposite sex on a cotton applicator and kept sealed until the presentation. Time spent sniffing the swab was quantitated with a stopwatch by a blinded human observer. Sniffing was scored when the nose was within 2 cm of the cotton swab. Each test session was conducted in a clean mouse cage containing fresh litter.

### Statistical analysis

Statistical analyses were performed using GraphPad Prism 6.01 software (Graph-Pad Software). Males and females were included and compared to determine if sex impacted our experimental results. All *n* values represent individual animals, unless stated otherwise (i.e. electrophysiology experiments). Notably, in analyses where multiple microglia were assayed per animal, all microglia analyzed for an individual animal were averaged to generate a single value per animal. When appropriate, animals were randomly assigned to conditions. Where possible, conditions were randomized to account for potential ordering effects. To ensure reproducibility, when relevant, experiments were performed at least three times independently. To avoid litter bias in the mouse experiments, experimental groups were composed of animals from different litters randomly distributed. All analyses were conducted with blinding to the experimental condition. Data are presented as scatter dot plot with bar, and the line at the mean ± S.E.M. (standard error of the mean). Comparisons between two groups following a normal distribution were analyzed using twotailed unpaired t-test with or without Welch’s correction, comparisons between two groups not following a normal distribution with Mann-Whitney test. Normality was assessed using the D’Agostino & Pearson omnibus normality test. When we compared cumulative distributions, we used the Kolmogorov-Smirnov test. When more than one variable was evaluated, we employed the two-way ANOVA with or without repeated measures (ANOVA and RM-ANOVA, respectively) with Sidak multiplicity-corrected post hoc comparisons to compare cohorts where appropriate. The statistical tests, including post-hoc tests for multiple comparisons, are reported in the figure legends along with the definition of *n*. For all analyses, α = 0.05.

## Acknowledgements

We thank Chris N. Parkhurst and Wenbiao B. Gan for providing the *Cx3cr1^creERT2^* knock-in mice, the *Cell* and *Tissue Imaging Facility* of the Institut du Fer à Moulin (namely Theano Eirinopoulou, Mythili Savariradjane and Xavier Marquès), where all image acquisitions and analyses have been performed, and the staff of the IFM animal facility (namely Baptiste Lecomte, Gaël Grannec, François Baudon, Anna-Sophia Mourenco, Emma Courteau and Eloise Marsan). We warmly thank Patricia Gaspar, Ludmilla Lokmane, Sonia Garel, Véronique Fabre and Jean Christophe Poncer for the discussion and revision. This work has been supported by grants from the Agence Nationale de la Recherche (ANR-17-CE16-0008, ANR-11-IDEX-0004-02), the Fondation pour la Recherche Médicale (Equipe FRM DEQ2014039529) and the Fédération pour la Recherche sur le Cerveau (FRC-2019-19F10).

## Author contributions

G.A., I.D., L.M. and A.Ro. designed the studies. G.A., I.D, A. Ro and L.M. wrote the manuscript, all authors revised it. G.A., I.D., F.E. and C.B. performed tamoxifen administration. G.A. performed immunofluorescence, image acquisition and analysis, Golgi-Cox staining and spine analyses. N.H. contributed to the design of DiOlistic labeling of dendritic spines, provided reagents and created the ImageJ Macro to analyze microglia/spines proximity. G.A. delivered DiI, performed image acquisition and analysis. C.L.M. contributed to the design and analysis of the electrophysiology experiments and provided reagents. G.A., M.D. and N.R.N. performed the electrophysiology experiments. A.Re. contributed to the design and analysis of the anterograde labeling of retinogeniculate projections and G.A. performed the intravitreal injections and image acquisition and analysis. I.D. performed behavioral experiments. G.A., I.D. and M.D. performed data analysis.

## Competing interests

The authors declare no competing interests.

---

**Fig. S1.**
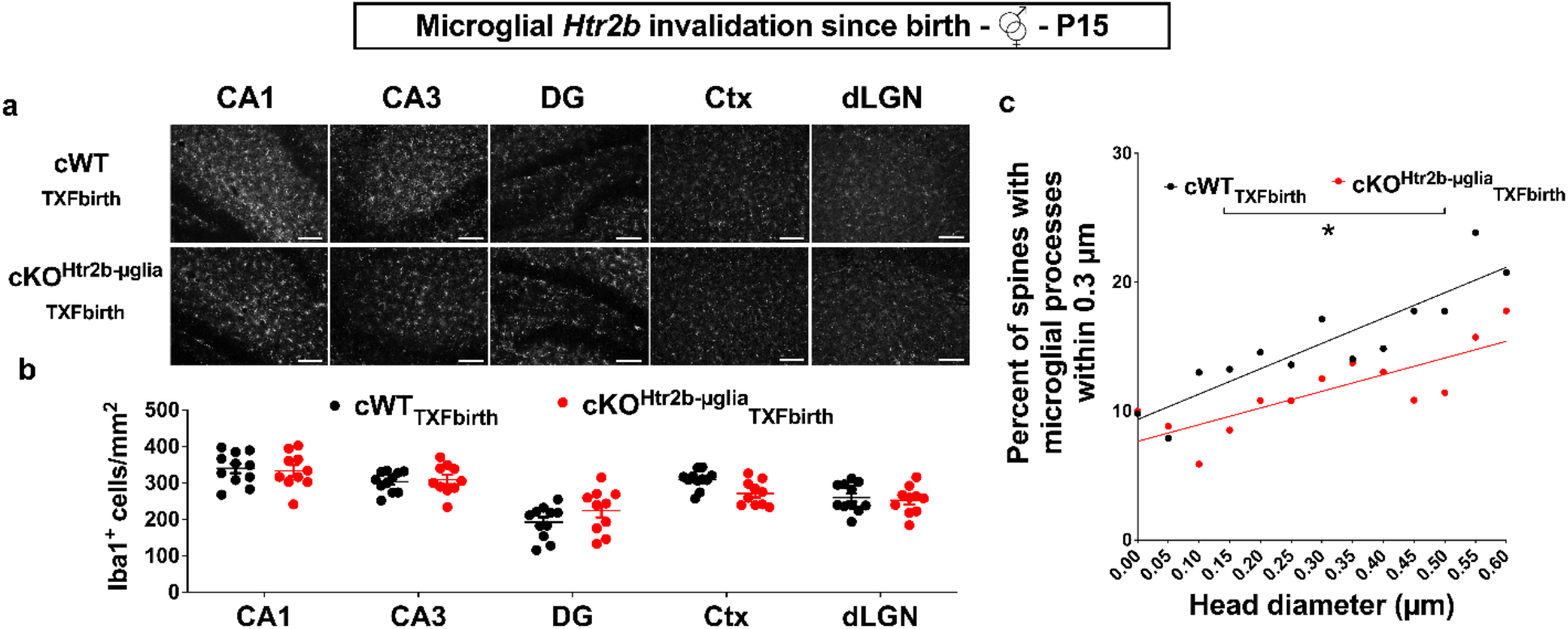
Early invalidation of *Htr2b* in microglia does not affect microglial density but decreases the percent of spines with microglial processes in proximity, independently of the spine head diameter. **a**, Representative images of Iba1^+^ cells on 20x images collected in several regions, including Cornu Ammonis (CA) 1, CA3 and dentate gyrus (DG) of the dorsal hippocampus, retrosplenial cortex (Ctx) and dorsal lateral geniculate nucleus of the thalamus (dLGN) in cWT_TXFbirth_ and cKO^Htr2b-μglia^_TXFbirth_ mice at P15. Scale bars: 100 μm. **b**, Densities of Iba1^+^ cells are similar in cWT_TXFbirth_ (black, *n* = 11 mice) and cKO^Htr2b-μglia^_TXFbirth_ (red, *n* = 10 mice) mice at P15. **c**, Absence of 5-HT signal in microglia since birth is associated to reduced percent of spines with microglial processes within 0.3 μm at P15, independently of the protrusion head diameter (cWT_TXFbirth_ mice, *n* = 13 mice; cKO^Htr2b-μglia^_TXFbirth_ mice, *n* = 12 mice; Kolmogorov-Smirnov test, P = 0.0461). Graphs show mean±s.e.m. and points represent individual animals. *p < 0.05.

**Fig. S2.**
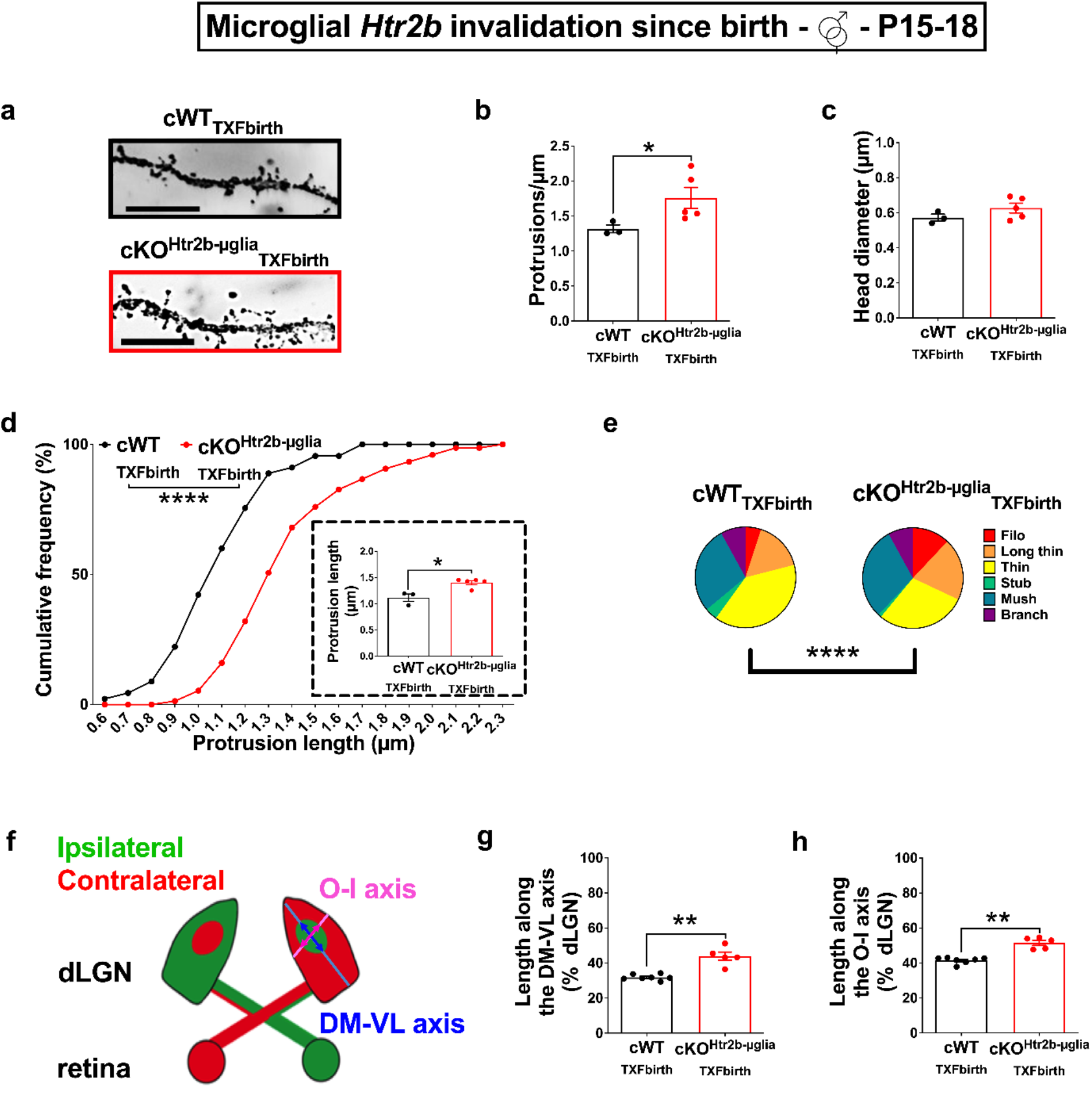
Early invalidation of *Htr2b* in microglia impacts on the maturation of the dLGN and on cortical spines. **a**, Representative images of apical secondary dendrites labeled with Golgi staining in L2/L3 principal neurons, scale bar: 10 μm. **b**, Quantification of protrusion density on secondary apical dendrites of L2/L3 principal neurons in cWT_TXFbirth_ and cKO^Htr2b-μglia^_TXFbirth_ mice at P15 obtained averaging 15 dendrites/mouse, that shows a significant increase in protrusion density in cKO^Htr2b-μglia^_TXFbirth_ mice (Mann-Whitney test, p = 0.0357). **c**, Similar average head diameter of protrusions on secondary apical dendrites of L2/L3 cortical neurons in cWT_TXFbirth_ and cKO^Htr2b-μglia^_TXFbirth_ mice at P15. **d**, Cumulative distribution plot for the average length of protrusions on secondary apical dendrites of L2/L3 principal neurons in cWT_TXFbirth_ and cKO^Htr2b-μglia^_TXFbirth_ mice at P15; note the significant rightward shift in cKO^Htr2b-μglia^_TXFbirth_ mice, as compared to cWT_TXFbirth_ mice (Kolmogorov-Smirnov test, p < 0.0001). In the inset, quantification of protrusion length obtained averaging 15 dendrites/mouse, that confirms a significant increase in average protrusion length in cKO^Htr2b-μglia^_TXFbirth_ mice (Mann-Whitney test, p = 0.0357). **e**, Pie charts showing the altered distribution of protrusions types in the cortex of cKO^Htr2b-μglia^_TXFbirth_ mice, as compared to cWT_TXFbirth_ mice at P15 (Chi-square test, p < 0.0001). **f**, A diagram of the dLGN showing the outer-inner (O-I) and dorsomedial-ventrolateral (DM-VL) axes. The lengths of the ipsilateral patch along the DM-VL axis (dark blue arrow) and the O-I axis (dark pink arrow) were divided by the lengths of the dLGN along the DM-VL axis (light blue line) and the O-I axis (light pink line), respectively. **g-h**, Extension of the ipsilateral retinal projections into the dLGN of cWT_TXFbirth_ and KO^Htr2b-μglia^_TXFbirth_ mice at P15. **g**, Ipsilateral retinal projections in cKO^Htr2b-μglia^_TXFbirth_ mice cover more territory along the DM-VL axis than in cWT_TXFbirth_ mice (Mann-Whitney test, p = 0.0025). **h**, Ipsilateral retinal projections in cKO^Htr2b-μglia^_TXFbirth_ mice cover more territory along the O-I axis than in cWT_TXFbirth_ mice (Mann-Whitney test, p = 0.0025). **b-c, d (inset), e**, cWT_TXFbirth_ mice, *n* = 3 mice, average of 15 dendrites per mouse; cKO^Htr2b-μglia^_TXFbirth_ mice, *n* = 5 mice, average of 15 dendrites per mouse; **d**, cWT_TXFbirth_ mice, *n* = 45 dendrites from 3 mice; cKO^Htr2b-μglia^_TXFbirth_ mice, *n* = 75 dendrites from 5 mice. **g-h**: cWT_TXFbirth_ mice, *n* = 7 mice; cKO^Htr2b-μglia^_TXFbirth_ mice, *n* = 5 mice. Graphs show mean±s.e.m. and points represent individual animals. *p < 0.05, **p < 0.01, ****p < 0.0001.

**Fig. S3.**
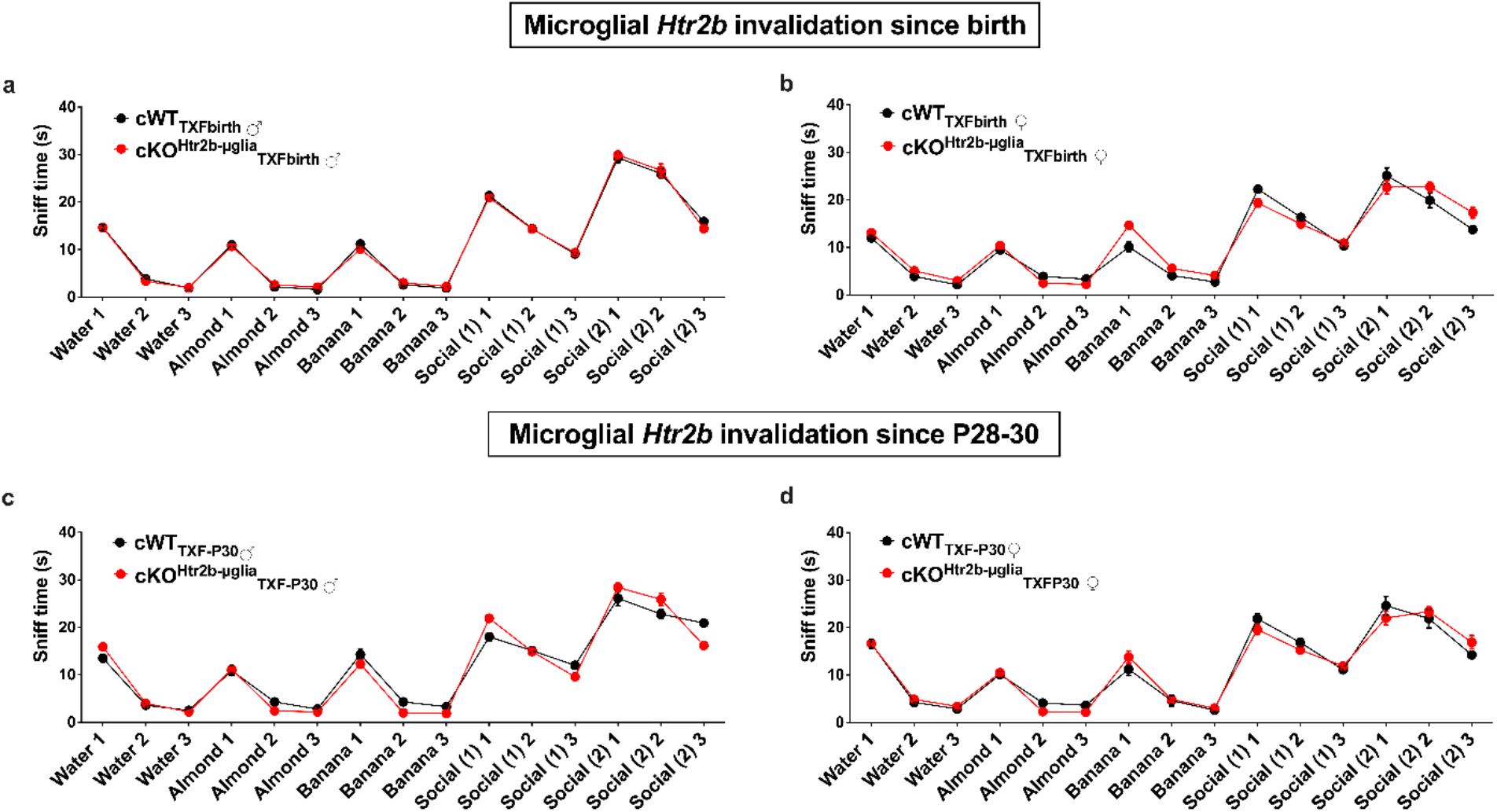
Early and late invalidations of *Htr2b* in microglia do not affect olfaction in mice. Olfactory habituation/dishabituation is identical in cWT_TXFbirth_ and cKO^Htr2b-μglia^_TXFbirth_ mice invalidated for the 5-HT2B receptors since birth (**a**, males; **b**, females) and since P28-30 (**c**, males; **d**, females). **a, b**: cWT_TXFbirth_, *n* = 12 mice; cKO^Htr2b-μglia^_TXFbirth_, *n* = 12 mice. **c**: cWT_TXFbirth_, *n* = 10 mice; cKO^Htr2b-μglia^_TXFbirth_, *n* = 10 mice. **d**: cWT_TXFbirth_, *n* = 10 mice; cKO^Htr2b-μglia^_TXFbirth_, *n* = 8 mice.

**Fig. S4.**
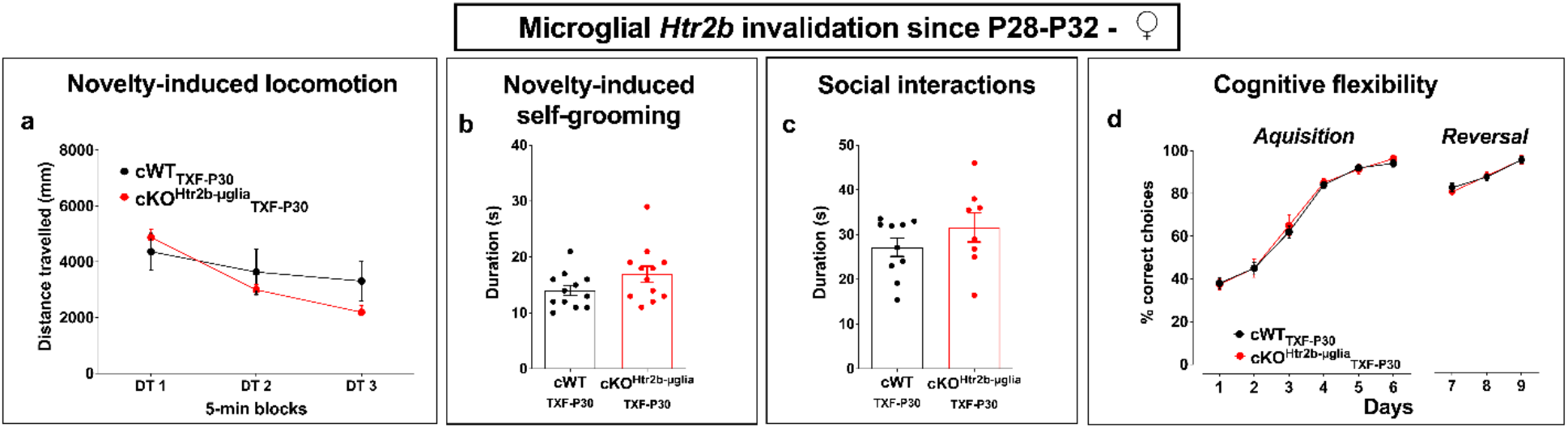
Late (P28-30) invalidation of microglial 5-HT_2B_ receptors does not affect activity in a novel environment, sociability nor flexibility in adult female mice. **a**, Similar distance travelled in response to a novel environment in 5-min blocks (DT) (two-way RM-ANOVA: significant main effect of DT *P* < 0.0001, *F*_2,36_ = 70.78). **b**, Similar time spent selfgrooming in response to a novel environment in cWTTXF-P30 and cKO^Htr2b-μglia^_TXF-P30_ adult female mice. **c**, Similar time spent interacting with a juvenile conspecific in a neutral territory in CWT_TXF-P30_ and cKO^Htr2b-μglia^_TXF-P30_ adult female mice. **d**, Y-maze reversal learning shows no difference in cognitive flexibility task in cWTTXF-P30 and cKO^Htr2b-μglia^_TXF-P30_ adult female mice. **a, c-d**: cWT_TXFbirth_, *n* = 10 female mice; cKO^Htr2b-μglia^_TXFbirth_, *n* = 8 female mice. **b**: cWT_TXFbirth_, *n* = 12 female mice; cKO^Htr2b-μglia^_TXFbirth_, *n* =12 female mice. Graphs show mean±s.e.m. and points represent individual animals.

## Notes

### Competing Interest Statement

The authors have declared no competing interest.

## References

1. Thion, M. S., Ginhoux, F. & Garel, S. Microglia and early brain development: An intimate journey. Science 362, 185–189 (2018).

2. Conio, B. et al. Opposite effects of dopamine and serotonin on resting-state networks: review and implications for psychiatric disorders. Mol Psychiatry 25, 82–93 (2020).

3. Lin, S.-H., Lee, L.-T. & Yang, Y. K. Serotonin and Mental Disorders: A Concise Review on Molecular Neuroimaging Evidence. Clin Psychopharmacol Neurosci 12, 196–202 (2014).

4. Marazziti, D. Understanding the role of serotonin in psychiatric diseases. F1000Res 6, 180 (2017).

5. Anderson, G. M., Horne, W. C., Chatterjee, D. & Cohen, D. J. The Hyperserotonemia of Autism. Ann NY Acad Sci 600, 331–340 (1990).

6. Migliarini, S., Pacini, G., Pelosi, B., Lunardi, G. & Pasqualetti, M. Lack of brain serotonin affects postnatal development and serotonergic neuronal circuitry formation. Mol Psychiatry 18, 1106–1118 (2013).

7. Witteveen, J. S. et al. Lack of serotonin reuptake during brain development alters rostral raphe-prefrontal network formation. Front. Cell. Neurosci. 7, (2013).

8. Teissier, A., Soiza-Reilly, M. & Gaspar, P. Refining the Role of 5-HT in Postnatal Development of Brain Circuits. Front. Cell. Neurosci. 11, 139 (2017).

9. Kolodziejczak, M. et al. Serotonin Modulates Developmental Microglia via 5-HT _2B_ Receptors: Potential Implication during Synaptic Refinement of Retinogeniculate Projections. ACS Chem. Neurosci. 6, 1219–1230 (2015).

10. Krabbe, G. et al. Activation of serotonin receptors promotes microglial injury-induced motility but attenuates phagocytic activity. Brain, Behavior, and Immunity 26, 419–428 (2012).

11. Béchade, C. et al. The serotonin 2B receptor is required in neonatal microglia to limit neuroinflammation and sickness behavior in adulthood. Glia 69, 638–654 (2021).

12. Etienne, F. et al. Two-photon Imaging of Microglial Processes’ Attraction Toward ATP or Serotonin in Acute Brain Slices. JoVE 58788 (2019) doi:10.3791/58788.

13. Schafer, D. P. et al. Microglia Sculpt Postnatal Neural Circuits in an Activity and Complement-Dependent Manner. Neuron 74, 691–705 (2012).

14. Upton, A. L. et al. Excess of Serotonin (5-HT) Alters the Segregation of Ispilateral and Contralateral Retinal Projections in Monoamine Oxidase A Knock-Out Mice: Possible Role of 5-HT Uptake in Retinal Ganglion Cells During Development. J. Neurosci. 19, 7007–7024 (1999).

15. Gaspar, P., Cases, O. & Maroteaux, L. The developmental role of serotonin: news from mouse molecular genetics. Nat Rev Neurosci 4, 1002–1012 (2003).

16. Pitychoutis, P. M., Belmer, A., Moutkine, I., Adrien, J. & Maroteaux, L. Mice Lacking the Serotonin Htr2B Receptor Gene Present an Antipsychotic-Sensitive Schizophrenic-Like Phenotype. Neuropsychopharmacol 40, 2764–2773 (2015).

17. Diaz, S. L. et al. 5-HT2B receptors are required for serotonin-selective antidepressant actions. Mol Psychiatry 17, 154–163 (2012).

18. Doly, S. et al. Serotonin 2B Receptors in Mesoaccumbens Dopamine Pathway Regulate Cocaine Responses. J. Neurosci. 37, 10372–10388 (2017).

19. Nikodemova, M. et al. Microglial numbers attain adult levels after undergoing a rapid decrease in cell number in the third postnatal week. Journal of Neuroimmunology 278, 280–288 (2015).

20. Dalmau, I., Vela, J. M., González, B., Finsen, B. & Castellano, B. Dynamics of microglia in the developing rat brain: Proliferation and Death of Microglia in Immature Brain. J. Comp. Neurol. 458, 144–157 (2003).

21. Weinhard, L. et al. Microglia remodel synapses by presynaptic trogocytosis and spine head filopodia induction. Nat Commun 9, 1228 (2018).

22. Vainchtein, I. D. et al. Astrocyte-derived interleukin-33 promotes microglial synapse engulfment and neural circuit development. Science 359, 1269–1273 (2018).

23. Stevens, B. et al. The Classical Complement Cascade Mediates CNS Synapse Elimination. Cell 131, 1164–1178 (2007).

24. Basilico, B. et al. Microglia shape presynaptic properties at developing glutamatergic synapses. Glia 67, 53–67 (2019).

25. Filipello, F. et al. The Microglial Innate Immune Receptor TREM2 Is Required for Synapse Elimination and Normal Brain Connectivity. Immunity 48, 979–991.e8 (2018).

26. Paolicelli, R. C. et al. Synaptic Pruning by Microglia Is Necessary for Normal Brain Development. Science 333, 1456–1458 (2011).

27. Zotova, E. et al. Inflammatory components in human Alzheimer’s disease and after active amyloid-β42 immunization. Brain 136, 2677–2696 (2013).

28. Weinhard, L. et al. Sexual dimorphism of microglia and synapses during mouse postnatal development: Sexual Dimorphism in Microglia and Synapses. Devel Neurobio 78, 618–626 (2018).

29. De Biase, L. M. et al. Local Cues Establish and Maintain Region-Specific Phenotypes of Basal Ganglia Microglia. Neuron 95, 341–356.e6 (2017).

30. Miyamoto, A. et al. Microglia contact induces synapse formation in developing somatosensory cortex. Nat Commun 7, 12540 (2016).

31. Risher, W. C., Ustunkaya, T., Singh Alvarado, J. & Eroglu, C. Rapid Golgi Analysis Method for Efficient and Unbiased Classification of Dendritic Spines. PLoS ONE 9, e107591 (2014).

32. Assali, A., Gaspar, P. & Rebsam, A. Activity dependent mechanisms of visual map formation--from retinal waves to molecular regulators. Semin Cell Dev Biol 35, 136–146 (2014).

33. D’Andrea, I., Alleva, E. & Branchi, I. Communal nesting, an early social enrichment, affects social competences but not learning and memory abilities at adulthood. Behavioural Brain Research 183, 60–66 (2007).

34. May, T., Adesina, I., McGillivray, J. & Rinehart, N. J. Sex differences in neurodevelopmental disorders. Current Opinion in Neurology 32, 622–626 (2019).

35. van den Berg, W. E., Lamballais, S. & Kushner, S. A. Sex-Specific Mechanism of Social Hierarchy in Mice. Neuropsychopharmacol 40, 1364–1372 (2015).

36. Zhan, Y. et al. Deficient neuron-microglia signaling results in impaired functional brain connectivity and social behavior. Nat Neurosci 17, 400–406 (2014).

37. Badimon, A. et al. Negative feedback control of neuronal activity by microglia. Nature 586, 417–423 (2020).

38. Assali, A. et al. RIM1/2 in retinal ganglion cells are required for the refinement of ipsilateral axons and eye-specific segregation. Sci Rep 7, 3236 (2017).

39. Thion, M. S. et al. Microbiome Influences Prenatal and Adult Microglia in a Sex-Specific Manner. Cell 172, 500–516.e16 (2018).

40. Villa, A. Sexual differentiation of microglia. Frontiers in Neuroendocrinology 9 (2019).

41. Hanamsagar, R. et al. Generation of a microglial developmental index in mice and in humans reveals a sex difference in maturation and immune reactivity: HANAMSAGAR et al. Glia 65, 1504–1520 (2017).

42. Jovanovic, H. et al. Sex differences in the serotonin 1A receptor and serotonin transporter binding in the human brain measured by PET. NeuroImage 39, 1408–1419 (2008).

43. Ohsawa, K. et al. P2Y _12_ receptor-mediated integrin-β1 activation regulates microglial process extension induced by ATP. Glia NA-NA (2010) doi:10.1002/glia.20963.

44. Sipe, G. O. et al. Microglial P2Y12 is necessary for synaptic plasticity in mouse visual cortex. Nat Commun 7, 10905 (2016).

45. Maroteaux, L., Béchade, C. & Roumier, A. Dimers of serotonin receptors: Impact on ligand affinity and signaling. Biochimie 161, 23–33 (2019).

46. von Kügelgen, I. & Hoffmann, K. Pharmacology and structure of P2Y receptors. Neuropharmacology 104, 50–61 (2016).

47. Oliver, K. H., Duvernay, M. T., Hamm, H. E. & Carneiro, A. M. D. Loss of Serotonin Transporter Function Alters ADP-mediated Glycoprotein αIIbβ3 Activation through Dysregulation of the 5-HT2A Receptor. Journal of Biological Chemistry 291, 20210–20219 (2016).

48. Baqi, Y. & Müller, C. E. Antithrombotic P2Y12 receptor antagonists: recent developments in drug discovery. Drug Discovery Today 24, 325–333 (2019).

49. Saini, H. K. et al. Therapeutic Potentials of Sarpogrelate in Cardiovascular Disease*. Cardiovascular Drug Reviews 22, 27–54 (2006).

50. Percie du Sert, N. et al. The ARRIVE guidelines 2019: updated guidelines for reporting animal research. http://biorxiv.org/lookup/doi/10.1101/703181 (2019) doi:10.1101/703181.

51. Parkhurst, C. N. et al. Microglia Promote Learning-Dependent Synapse Formation through Brain-Derived Neurotrophic Factor. Cell 155, 1596–1609 (2013).

52. Belmer, A. et al. Positive regulation of raphe serotonin neurons by serotonin 2B receptors. Neuropsychopharmacol 43, 1623–1632 (2018).

53. Pitulescu, M. E., Schmidt, I., Benedito, R. & Adams, R. H. Inducible gene targeting in the neonatal vasculature and analysis of retinal angiogenesis in mice. Nat Protoc 5, 1518–1534 (2010).

54. Goldmann, T. et al. A new type of microglia gene targeting shows TAK1 to be pivotal in CNS autoimmune inflammation. Nat Neurosci 16, 1618–1626 (2013).

55. Peng, J. et al. Microglia and monocytes synergistically promote the transition from acute to chronic pain after nerve injury. Nat Commun 7, 12029 (2016).

56. Wolf, Y., Yona, S., Kim, K.-W. & Jung, S. Microglia, seen from the CX3CR1 angle. Front. Cell. Neurosci. 7, (2013).

57. Yona, S. et al. Fate Mapping Reveals Origins and Dynamics of Monocytes and Tissue Macrophages under Homeostasis. Immunity 38, 79–91 (2013).

58. Franklin, K. B. J. & Paxinos, G. Paxino’s and Franklin’s the Mouse Brain in Stereotaxic Coordinates: Compact 5th Edition. (Elsevier Science & Technology, 2019).

59. Schafer, D. P., Lehrman, E. K., Heller, C. T. & Stevens, B. An Engulfment Assay: A Protocol to Assess Interactions Between CNS Phagocytes and Neurons. JoVE 51482 (2014) doi:10.3791/51482.

60. Young, K. & Morrison, H. Quantifying Microglia Morphology from Photomicrographs of Immunohistochemistry Prepared Tissue Using ImageJ. JoVE 57648 (2018) doi:10.3791/57648.

61. Heck, N., Betuing, S., Vanhoutte, P. & Caboche, J. A deconvolution method to improve automated 3D-analysis of dendritic spines: application to a mouse model of Huntington’s disease. Brain Struct Funct 217, 421–434 (2012).

62. Rodriguez, A., Ehlenberger, D. B., Dickstein, D. L., Hof, P. R. & Wearne, S. L. Automated Three-Dimensional Detection and Shape Classification of Dendritic Spines from Fluorescence Microscopy Images. PLoS ONE 3, e1997 (2008).

63. Giralt, A. et al. Pyk2 modulates hippocampal excitatory synapses and contributes to cognitive deficits in a Huntington’s disease model. Nat Commun 8, 15592 (2017).

64. Zaqout, S. & Kaindl, A. M. Golgi-Cox Staining Step by Step. Front. Neuroanat. 10, (2016).

65. Rebsam, A., Petros, T. J. & Mason, C. A. Switching Retinogeniculate Axon Laterality Leads to Normal Targeting but Abnormal Eye-Specific Segregation That Is Activity Dependent. Journal of Neuroscience 29, 14855–14863 (2009).

66. Hayakawa, I. & Kawasaki, H. Rearrangement of Retinogeniculate Projection Patterns after Eye-Specific Segregation in Mice. PLoS ONE 5, e11001 (2010).

67. Silverman, J. L., Tolu, S. S., Barkan, C. L. & Crawley, J. N. Repetitive Self-Grooming Behavior in the BTBR Mouse Model of Autism is Blocked by the mGluR5 Antagonist MPEP. Neuropsychopharmacol 35, 976–989 (2010).

68. Scattoni, M. L., Gandhy, S. U., Ricceri, L. & Crawley, J. N. Unusual Repertoire of Vocalizations in the BTBR T+tf/J Mouse Model of Autism. PLoS ONE 3, e3067 (2008).

69. D’Andrea, I. et al. Lack of kinase-independent activity of PI3Kγ in locus coeruleus induces ADHD symptoms through increased CREB signaling. EMBO Mol Med 7, 904–917 (2015).

70. Dandrea, I., Alleva, E. & Branchi, I. Communal nesting, an early social enrichment, affects social competences but not learning and memory abilities at adulthood. Behavioural Brain Research 183, 60–66 (2007).

71. Moy, S. et al. Mouse behavioral tasks relevant to autism: Phenotypes of 10 inbred strains. Behavioural Brain Research 176, 4–20 (2007).

72. Van der Borght, K., Havekes, R., Bos, T., Eggen, B. J. L. & Van der Zee, E. A. Exercise improves memory acquisition and retrieval in the Y-maze task: Relationship with hippocampal neurogenesis. Behavioral Neuroscience 121, 324–334 (2007).

73. Yang, M. & Crawley, J. N. Simple Behavioral Assessment of Mouse Olfaction. Current Protocols in Neuroscience 48, (2009).

